# The more the merrier? Multivariate approaches to genome-wide association analysis

**DOI:** 10.1101/610287

**Authors:** César-Reyer Vroom, Christiaan de Leeuw, Danielle Posthuma, Conor V. Dolan, Sophie van der Sluis

## Abstract

The vast majority of genome-wide association (GWA) studies analyze a single trait while large-scale multivariate data sets are available. As complex traits are highly polygenic, and pleiotropy seems ubiquitous, it is essential to determine when multivariate association tests (MATs) outperform univariate approaches in terms of power. We discuss the statistical background of 19 MATs and give an overview of their statistical properties. We address the Type I error rates of these MATs and demonstrate which factors can cause bias. Finally, we examine, compare, and discuss the power of these MATs, varying the number of traits, the correlational pattern between the traits, the number of affected traits, and the sign of the genetic effects. Our results demonstrate under which circumstances specific MATs perform most optimal. Through sharing of flexible simulation scripts, we facilitate a standard framework for comparing Type I error rate and power of new MATs to that of existing ones.

## Introduction

Genome-wide association (GWA) studies aim to identify single nucleotide polymorphisms (SNPs) that are associated with (i.e., explain variation in) continuous traits (e.g., height, blood pressure, BMI), or in the liability underlying dichotomous (disease) traits (e.g., schizophrenia, cancer, Type II diabetes). Most GWA studies are univariate in the sense that they focus on a single trait. However, often data on multiple correlated traits are available and sometimes traits treated as univariate are actually multivariate in nature. For instance, GWA studies on metabolic syndrome (e.g., Zhu et al., 2017, Kristiansson et al., 2012) base the case-control status on the joint evaluation of multiple measures (e.g., waist circumference, body mass index, blood pressure, and various blood measures). Similarly, GWA studies on psychiatric disorders like major depressive disorder (e.g., Howard et al, 2018, Wray et al., 2018) generally use case-control status variables that originate in the joint evaluation of multiple clinical criteria, and GWA studies on cognitive ability use cognitive scores that summarize the performance on batteries of cognitive tests covering e.g., vocabulary, general knowledge, and memory (e.g., Savage et al., 2018, Benyamin et al., 2014; Davis et al., 2010).

With increasing availability of multivariate information (e.g., UK Biobank), and knowing that pleiotropy is wide-spread both within and between trait domains (Watanabe et al., in revision), it is important to determine the circumstances in which a multivariate approach has greater statistical power than the standard univariate test to detect an associated SNP, which we henceforth will generally refer to as the genetic variants (GV, plural GVs). As GWA studies use a stringent correction for multiple testing (usually α is set to 5 × 10^−8^, Pe’er et al., 2008, Sham & Purcell, 2014), and effect sizes of individual GVs are expected to be small (e.g. Visscher et al., 2012, 2017; Psychiatric GWAS Consortium, 2009), statistical power remains a pivotal concern in GWA studies, despite increasing sample sizes. Besides increasing study sample sizes, exploiting the multivariate nature of GWA data sets may under some circumstances, as we will demonstrate here, increase the *statistical power* to detect GVs.

Numerous multivariate association tests (MATs) are available. We define a MAT as any test that formalizes the statistical association between a GV and a set of *m* traits that are measured in the same individual. MATs differ in several respects, such as their ability to accommodate missing values or traits of different measurement levels (e.g., a mix of continuous and dichotomous traits). The power of MATs has been subject of investigation, but the scope of the settings in which power was studied was generally limited: simulation scenarios often featured just a few (e.g., 2 or 3; He et al., 2013, Wu & Pankow, 2015), uniformly correlated traits, only GVs that affect all traits in the analysis (Galesloot et al., 2014; Van der Sluis et al., 2013; Aschard et al., 2014, Suo et al., 2013; Yang et al., 2016), or only same-sign GV effects (e.g., Porter & O’Reilly, 2017). Reality is, however, often more complex, and the *true genotype-phenotype* model (i.e., the model describing the relations between the *m* traits and the GV as they are in reality) is usually unknown. To determine the circumstances in which MATs perform best in terms of power the following should be considered: the number of traits in the simulations, the correlational patterns between the traits (e.g., both uniform and block-wise), the generality of GV effects (i.e., the number of traits affected by the GVs), and the sign of the GV effects (i.e., allowing the reality of opposite effects).

The aim of this Review is to provide a classification of available MATs, to give an overview of their defining characteristics, to inspect their *Type I error rate*, and to compare their statistical power to detect GVs under a multitude of realistic circumstances. We classify MATs based on the underlying statistical model, and explicate their associated hypotheses. We inspect Type I error rates in various circumstances, given various values of *criterion level α*, and we identify the circumstances in which conducting multivariate analyses is (dis)advantageous in terms of statistical power. We do so through extensive simulation in which we investigate the effects of the factors mentioned above: the number of traits in the analysis, the correlational pattern between the traits, and generality and sign of the GV effects. We show that the power of MATs can vary considerably as a function of the true genotype-phenotype model (e.g., in consequence of the presence of unaffected traits or opposite GV-effects). Overall, these results facilitate the choice of the most appropriate and optimal MATs in future multivariate GWA studies. Through sharing of flexible simulation scripts (https://ctg.cncr.nl/software/), we facilitate prospective application of a standard verification framework within which the statistical power and Type I error rate of new MATs can be compared to that of existing ones.

## 1. Classification of MATs

A wide range of MATs are available (see Table 1 for an overview of the MATs included in this paper). Following Yang and Wang’s conceptual classification (Yang & Wang, 2012), we distinguish transformation-based MATs, regression-based MATs, and combination tests. We discuss each class of MATs and provide a short statistical description of the MATs included in this review in Boxes 1–3. These descriptions provide a basic understanding of the statistical properties of individual MATs, which furthers insight into their specific strengths and weaknesses. For a non-statistical overview of all included MATs, we refer to Table 1. Note that in each MAT, the predictor of main interest is a single genetic variant, i.e., a potential GV. In practice, however, additional predictors (i.e., covariates) are standardly included in the model such as the age and sex of participants, and genetic principle components (obtained using e.g. Eigenstrat (Price et al., 2006) or FlashPC2 (Abraham & Inouye, 2014)) to correct for population stratification.

**Table 1.**
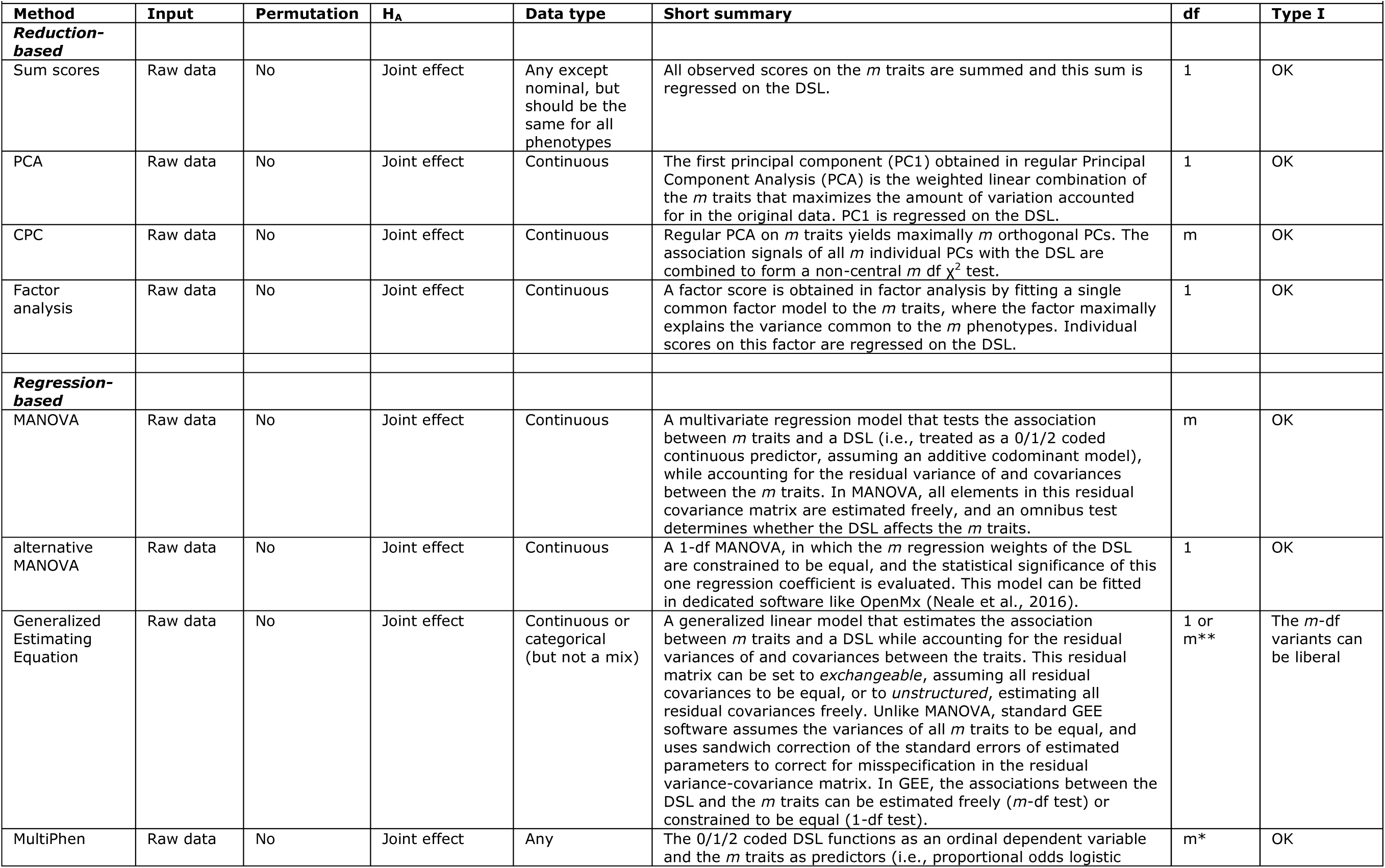
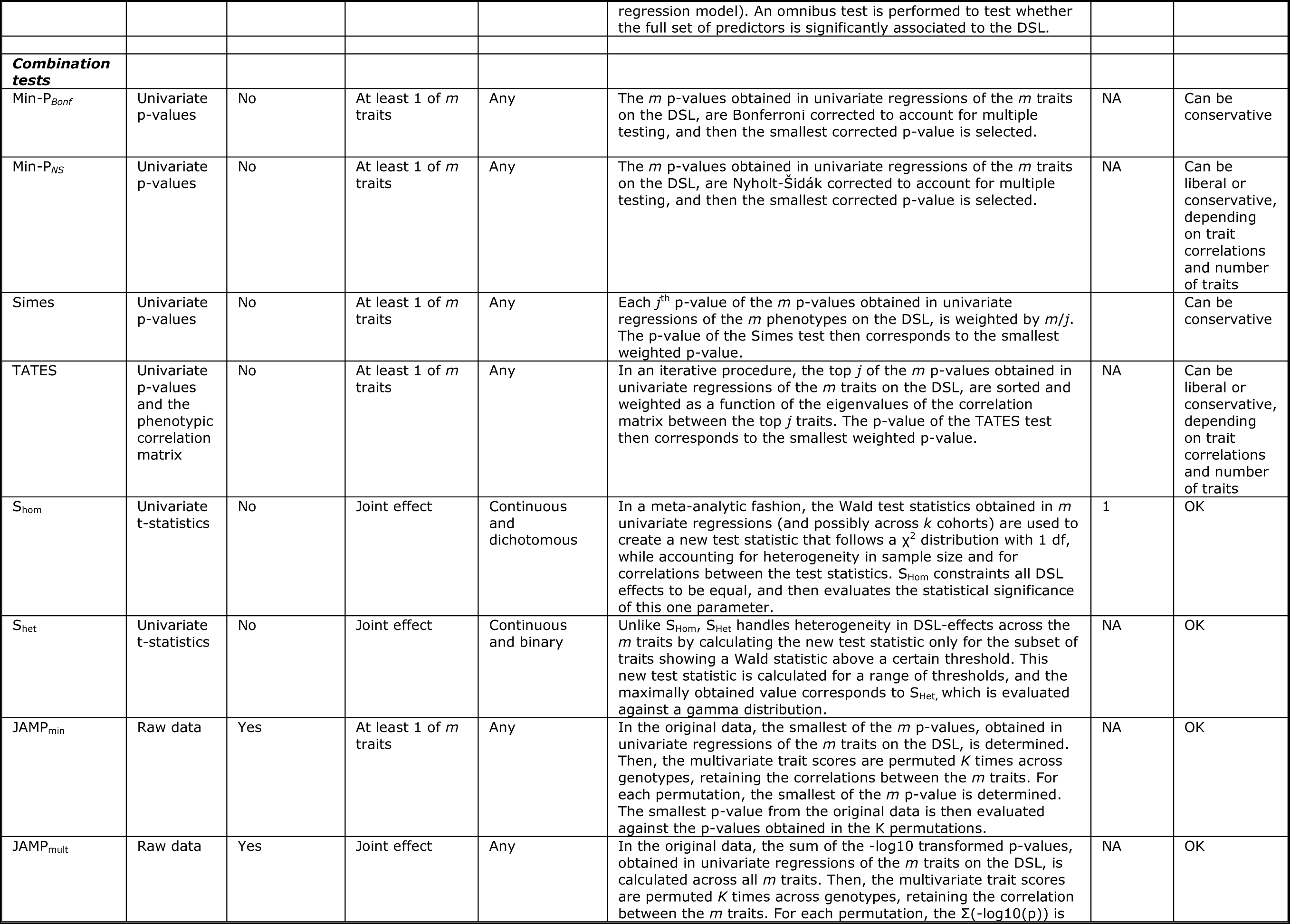
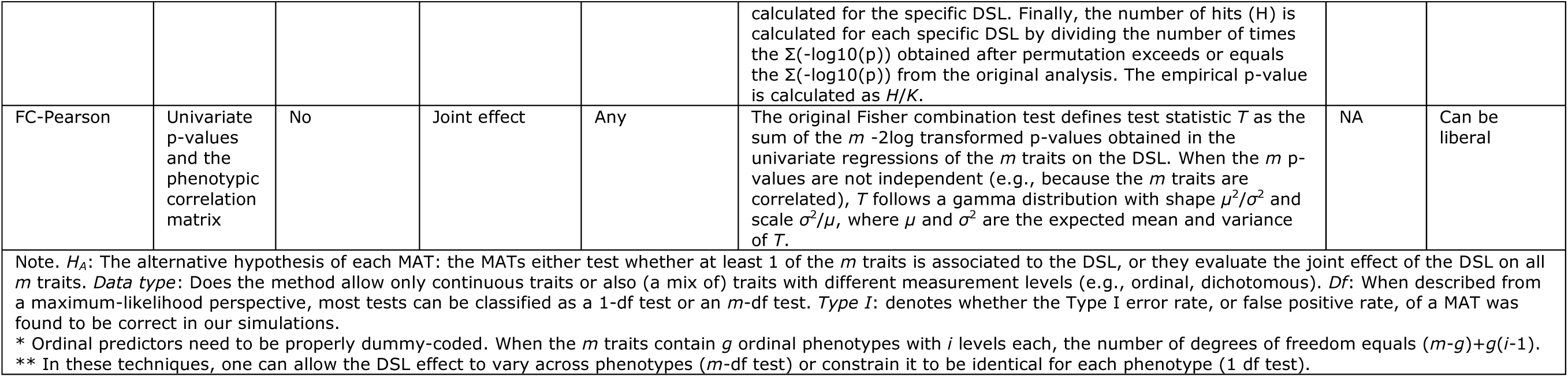
Classification of multivariate methods.

### Box 1 Transformation-based techniques

#### Sum-scores

In psychology and psychiatry, sum scores are often used to summarize multivariate responses to items on tests (e.g., cognitive ability), questionnaires (e.g., personality), and clinical instruments and interviews (e.g., depression). In psychiatric studies, the sum-score is often dichotomized to obtain a binary case-control status variable, although this may lower the power to detect a possible GV (e.g. Van der Sluis et al., 2012; Lee & Wray, 2013). In the case of an unweighted sum score (i.e., b_1_ to b_m_ in Eq 1 are set to one), the variance of a sum score equals the sum of all entries of the *m* × *m* variance-covariance matrix of the *m* traits. How well the GV can be detected through the sum score thus not only depends on the effect size of the GV, but also on the number of traits it affects. The contribution of *global* GVs, i.e., GVs that affect all or multiple of the *m* traits, to the variance of the sum is generally larger than to the variance of the underlying elements, so that the power to detect global GVs can benefit from using a sum-score. In contrast, GVs that affect only 1 or a few of the traits (i.e., *local* variants) contribute relatively little to the variance of the sum. Importantly, however: how well a sum-score reflects the GV-effect(s) also depends on the magnitude of the variances and covariances conditional on the GV: if these conditional (co)variances are relatively small, then the *signal-to-noise ratio* will be better than if the conditional (co)variances are large (see Supplemental Information for a more formal discussion of this topic).

#### Principal Component Analysis (PCA)

PCA is used to transform a set of *m* correlated standardized traits into a set of maximally *m* orthogonal (i.e., uncorrelated) linear combinations of these traits, the new variates being denoted as Principal Components (PC). For the first PC (PC1), the weights *b*_1_…*b*_m_ in Eq. 1 are chosen such that the variance of PC1 is maximized. If the correlations between the *m* traits are equal (i.e., homogeneous), then PC1 will correlate 1 with the sumscore (as, b_1_ = b_2_ =…=b_m_). PC1 provides a summary of the full set of *m* traits. Additional PCs may be considered if the variance of PC1 is judged to be too small. In the psychometric context, where the *m* traits are generally items measuring a given latent trait (e.g., neuroticism), PC1 is viewed as a proxy of that latent trait. Assuming that PCA was used to reduce multivariate information, we focus on the analysis of PC1 (see Supplemental Information).

#### Combined PC test (CPC test)

As PCA is conducted on the trait information and does not involve genetic information, of all PCs obtainable in PCA of a set of *m* traits, PC1 does not necessarily have the strongest association with the GV. In PCA’s iterative procedure, the variance in *y*_1_…*y*_m_ that is not accounted for by preceding PCs, can be accounted for by successive PCs. The weights of successive PCs are chosen such that again their variance is maximized and that they are uncorrelated with preceding PCs. Capitalizing on the fact that the *m* extracted PCs are uncorrelated (orthogonal), the combined PC test (CPC test) evaluates the association of the GV to all *m* PCs simultaneously by reference to a *χ*^2^-distribution with *m* degrees of freedom (Aschard et al., 2014).

#### Common factor analysis

As a data transformation method, factor analysis resembles PCA: just like one may use PC1, one can also fit a single common factor model to the *m* traits, calculate the scores on the common factor (i.e., factor scores), and use this factor score as dependent variable in GWA studies. In the single common factor model, the weights *b*_1_…*b*_m_ in Eq. 1 are chosen such that the variance explained by the new variate ỹ in the set of *m* traits is maximized, i.e., ỹ maximally represent the variance common to the *m* traits. While PCA concerns the total variance of the traits, factor analysis thus focusses on the covariance shared by the *m* traits (also denoted as ‘communality’). This common factor obtained in factor analysis may be viewed as a substantive variable: a common cause of (and as such a source of covariance among) the *m* traits (Lawley & Maxwell, 1971). For instance, the covariance between *m* neuroticism symptoms is assumed to originate in the fact that all *m* symptoms are caused by the underlying latent trait “neuroticism”. PCA and factor analysis are thus conceptually different: PCA components are merely statistically optimal linear variates, while the factors in factor analysis are often assumed to actually represent a theoretical construct (e.g., neuroticism). In addition, the residuals of the *m* traits, i.e., the unique parts of *y*_1_…*y*_m_ that are not explained by the variate ỹ, are assumed to be uncorrelated in factor analysis, while no such assumption is made in PCA. In practice, however, PCA and factor analysis often yield very similar result, e.g. when the communality of the traits is high (i.e., the variance shared by the *m* traits is high compared to the unique variance of the traits). Assuming that factor analysis was used to reduce multivariate information, we focus on the analysis of factor scores obtained in a single common factor model (see Supplemental Information).

#### Canonical Correlation Analysis

Canonical Correlation Analysis (CCA) extracts for each GV under study the linear combination of *m* traits (i.e., variate) that explains the largest amount of covariance with that specific GV (Solovieff et al., 2013). The weights of the new variate thus differ between GV, and reveal which traits are the most strongly associated to a specific GV. CCA is thus the only transformation-based technique that uses the information from the GV to create the new variate. CCA is implemented in the widely used GWA package PLINK (Ferreira & Purcell, 2009). However, assuming an *additive codominant genetic model* in which the GV, coded 0/1/2 for the number of minor alleles, is treated as a continuous predictor (i.e., a “covariate”, rather than a “factor”), CCA is known to perform identically to MANOVA and therefore does not feature as a separate MAT in our study.

### Box 2 Regression-based techniques

All regression-based techniques described here assume that conditional on the effect of the GV, the data of the *m* traits follow a multivariate normal distribution.

#### MANOVA

In standard MANOVA, the *m* × *m* symmetrical background covariance matrix **Σ_E_** is unconstrained, i.e., it has ((*m*+1)\**m*)/2 freely estimated elements (covariances and variances). In terms of a likelihood ratio test (asymptotically equal to the F-test used to evaluate MANOVA), standard MANOVA is an *m*-df omnibus test of the null hypothesis that the *m* regression coefficients are all zero (no association). For comparison, we also ran simulations for a 1-df MANOVA (fitted in the R package OpenMx (Neale et al., 2016), in which the *m* regression weights of the GV are constrained to be equal, and the null-hypothesis is that this regression coefficient is zero (no association).

#### Generalized Estimating Equations (GEE)

In GEE, one can specify various structures for **Σ_E_**, which is modeled as **Δ_E_P_E_Δ_E_**, where **P_E_** is the residual correlation matrix between the *m* traits conditional on all predictors in the model, and **Δ_E_** is a diagonal matrix with the *m* residual standard deviations of the *m* traits constrained to be equal. In GEE, the structure of correlation matrix **P_E_**, i.e., the working correlation matrix, is user-specified. In order of parsimony, plausible choices for **P_E_** are “independent” (**P_E_ =I;** the *m* traits show no correlation conditional on the GV), “exchangeable” (all conditional correlations between the *m* traits are equal), and “unstructured” (i.e., all conditional correlation are freely estimated).

Standard GEE software uses sandwich correction of the standard errors of estimated parameters to correct for the possible misspecification of **Σ_E_** (ref Dobson). As demonstrated elsewhere (e.g., Minica et al. 2015), the degree of misspecification does have a bearing on the power of the sandwich corrected test. In our simulations, we specified 1-df versions of ‘exchangeable’ and ‘unstructured’ GEE models (i.e., the *m* regression weights of the modelled GV were constrained to be identical). As *m*-df versions of ‘exchangeable’ and ‘unstructured’ GEE models yield identical results (see Supplemental Information), we only included the results of GEE-unstructured *m*-df models in our main discussion, but results for the GEE ‘exchangeable’ *m*-df model are available in the Tables S7-S12.

#### Linear Mixed Models (LMM)

Linear mixed effects models are an extension of the multivariate regression model, in which fixed effects are used to estimate the effects of the GV, and additional random effects account for the correlations among the *m* phenotypes (see e.g., Yang & Wang, 2012). In the genetics literature, LMM are frequently employed to model population substructure and relatedness in a univariate settings (e.g., EMMAX, GenABEL, FaST-LMM, Mendel, GEMMA and MMM, see Euahsunthornwattan et al (2014) for comparisons, and Yang et al (2014) for a discussion of potential pitfalls), but LMM can also be used to model e.g. multivariate gene-environment interaction (Moore et al., 2018) or to accommodate multivariate data (e.g., Zhou & Stephens, 2014). In principle, LMM can handle multiple sources of clustering or correlation (e.g., multivariate data and familial relatedness or population substructure simultaneously). Because LMM often failed to converge in our simulations (especially with larger *m*), and Type I error rates were severely off for the *m*-df variant, we excluded LMM from our main discussion, but all results are available in the Tables S7-S12.

#### Multiphen: reversed ordinal multiple regression

The MultiPhen procedure (O’Reilly et al., 2012) reverses the regression model by treating the GV as an ordinal dependent variable, and the *m* traits as predictors. This has the practical advantage of rendering distributional assumption concerning the phenotypes (e.g., conditional multivariate normality, see Table 1) unnecessary; the *m* phenotypes can be a mix of continuous and categorical (appropriately dummy-coded) variables. The procedure is implemented in an R-package (‘MultiPhen’). MultiPhen tests the *m* df null-hypothesis that the *m* regression coefficients are zero.

### Box 3 Combination tests

#### Minimal p-values: min-PNS and min-PBonf

Minimal p-value approaches use the *m* p-values obtained in univariate analyses, correct these p-values for multiple testing, and then select the smallest. Specifically, to obtain the Bonferroni-corrected minimal p-value, min-P_Bonf_, first all original p-values are multiplied by *m* to obtain the Bonferroni-corrected p-values, and then the minimal Bonferroni-corrected p-value is selected (Simes, 1986). To obtain the Nyholt-Šidák corrected minimal p-value, min-P_NS_ (O’Reilly et al., 2012), one first establishes the *effective number* of traits *m*_e_, and this effective number of traits is then used to calculate the Sidak-corrected p-values as (1 − (1 − *p_org_*))*^m_e_^*. Nyholt (2004) proposed to calculate *m*_e_ as a function of the variance of all eigen values, which can be derived from the correlation matrix between the *m* traits.

#### Simes

To obtain the p-value for the original Simes test (Simes, 1986), *P*_S_, the *m* p-values obtained in *m* univariates association tests are first sorted ascendingly. Subsequently, each *j*th p-value (*j* running from 1 to *m*) is weighted with *m*/*j*, such that the lowest p-value is weighted with the largest weight (i.e., *m*/1) and the highest p-value is weighted with the smallest weight (i.e., *m*/*m*=1). The Simes p-value then corresponds to the smallest weighted p-value, i.e., 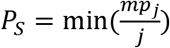

#### TATES: adjusted Simes test

As the original Simes test is conservative (Simes, 1986), and becomes more so with increasing correlations and increasing *m* (van der Sluis et al., 2018), Van der Sluis et al (2012) developed an adjusted Simes procedure denoted TATES (Trait-based Association Test that uses Extended Simes: based on Li et al., 2011). TATES weights in a fashion similar to Simes, except that the observed number of p-values *m* and *j* are replaced with the effective number of p-values *m*_e_ and *m*_ej_. Specifically, the TATES p-value *P_T_* is obtained as 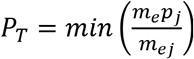, where *m_e_* denotes the effective number of independent p-values, and *m*_ej_ the effective number of p-values among the top *j* p-values. The effective number of p-values *m*_e_ and *m*_ej_ is established from eigenvalue decomposition of the correlation matrix between the *m* p-values, which can be approximated from the correlation matrix between the *m* traits (see Van der Sluis et al., 2012, 2018).

#### JAMP

The permutation-based software tool JAMP (Joint genetic Association of Multivariate Phenotypes, https://ctg.cncr.nl/software/jamp) incorporates two different multivariate tests: one that tests whether at least one of the *m* traits is associated to the GV (JAMP_min_), and one that assesses the joint association signal of the *m* traits to the GV (JAMP_mult_)^2^. Specifically, to calculate the empirical p-value for multivariate association, JAMP_mult_ uses permutation to control the Type I error rate and to adjust for correlations between the *m* traits. First, the univariate associations between the *m* traits and a GV are evaluated, and the GV-specific statistic *G*_o_ is calculated as 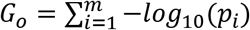, aggregating the signal across the *m* traits. Second, the *m* traits scores are permuted *J* times across the GV, keeping the correlations between the *m* traits intact. For each permutation, 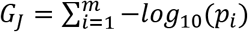 is calculated for the specific GV. Finally, the number of hits (*H*) is calculated for each GV by dividing the number of times *G*_J_ obtained on permuted data exceeds or equals *G*_o_ obtained on the original data. The empirical p-value (*P*_mult_) is then calculated as *P_mult_* = *H/J*.

In contrast, JAMP_min_ produces an empirical p-value (*P*_min_) associated with the hypothesis that at least one of the *m* traits is significantly associated with the GV. For each GV, the smallest of the *m* univariate p-values obtained in the original data is evaluated against the smallest of *m* univariate p-values obtained in each of the *J* permutations. In our simulations, the number of permutations *J* was set to 1000.

S_Hom_. In a meta-analytic fashion, S_Hom_ (Zhu et al., 2015) uses the Wald test statistics obtained in *m* univariate GWASs (and possibly across *k* cohorts) to create a new test statistic that follows a χ^2^ distribution with 1 df. S_Hom_ accounts for heterogeneity in sample size and for correlations between the test statistics. As a 1 df test, S_Hom_ constraints all GV effects to be the same, and then tests the omnibus hypothesis that this 1 GV-parameter is 0. S_Hom_ is thus most powerful when the GV effects are homogeneous in size and sign across the *m* traits.

S_Het_. S_Het_ is equivalent to S_Hom_ but specifically handles heterogeneity in GV-effects across the *m* traits by calculating the new test statistic only for the subset of traits showing a Wald statistic above a certain threshold. This new test statistic is calculated for a range of thresholds, and the maximally obtained value corresponds to S_Het_. The significance of S_Het_ is obtained through simulation of a Gamma distribution (see Supplemental Information for details). Like S_Hom_, S_Het_ tests the omnibus hypothesis that all included effects are zero. Because of the selection, S_Het_ is expected to be more powerful than S_Hom_ when the GV-effects are heterogeneous in size and/or sign across the *m* traits.

#### FC-Pearson test: adjusted Fisher Combination test

Let *p*_1_…*p*_m_ be the p-values obtained in the univariate regressions of the *m* traits on a GV. The original FC-test is calculated as 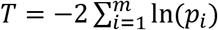 (Fisher, 1932). If the *m* traits are uncorrelated, the original FC test statistic *T* is chi-squared distributed with 2*m* dfs. However, if the *m* traits are correlated, this original test has highly inflated Type I error rate (Fisher, 1932; van der Sluis et al., 2012). For *m* correlated traits, it can be shown (Brown and Yang, ref 27/28 in Yang et al, 2016) that, under the null hypothesis of no association between the GV and the *m* traits, *T* follows a scaled chi-squared distribution, or equivalently a specific gamma distribution with shape parameter that can be derived from the mean (µ) and variance (σ^2^) of test statistic T. Yang et al. (2016) established an approximation of µ and σ in case of *m* continuous correlated traits. Just like the original FC-test, this adjusted test, referred to as the FC-Pearson test, tests the hypothesis that the aggregated GV-signal present in the set of *m* traits deviates significantly from 0.

### Transformation-based MATs

The simplest way to deal with a multivariate problem is by reducing it to a univariate problem through transformation of the multivariate information. Given *N* subjects and *m* traits *y*_1_…*y*_m_, a single new *variate ỹ* for subject *i* can be created that is a linear combination of these *m* traits:

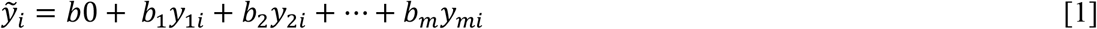

where the weights *b*_1_…*b*_m_ determine how much each original trait contributes to the new variate. All transformation-based MATs are aimed at variable reduction. The following transformation-based MATs are included in this review and their characteristics (e.g., how the weights *b*_1_…*b*_m_ in Eq 1 are determined) are described in Box 1: sum-score analysis, Principal Component Analysis (PCA), the Combined Principal Components test (CPC, Asschard et al., 2014), and common factor analysis. Important to note is that all transformation-based MATs determine the weights in Eq. 1 independently of the association of the *m* traits with the GV (e.g., in factor analysis, the weights depend on the correlations among the *m* phenotypes only). That is, all transformation-based MATs first transform the data solely based on the phenotypic information, and only then consider the possible association of this new variate with the GV (generally using a univariate regression model).

### Regression-based MATs

In a multivariate GWA settings, one focusses on the association between a set of *k* predictors (the GV and the covariates), and a set of *m* traits. Given *N* subject, *m* traits and *k* predictors, this multivariate (referring to the number of dependent variables) multiple (referring to the number of predictors) regression model can be represented as:

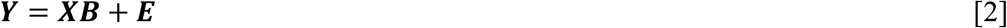

Here, **Y** is the N × *m* matrix of trait scores. **X** is the design matrix, i.e., a N × (*k*+1) matrix of predictor scores in which the first column usually is a unit vector that serves to estimate the *m* trait-specific intercepts. **B** is a (*k*+1) × *m* matrix of regression weights with the first row containing *m* trait-specific intercepts, and the subsequent *k* rows containing *m* trait-specific regression weights for the *k* predictors. The *m* regression weights on the row corresponding to the GV are usually all freely estimated, giving rise to an *m* degrees of freedom (df) omnibus test (i.e., the GV is allowed to affect the *m* traits differently). The *m* weights may be constrained to be equal, thus giving rise to a 1-df test (i.e., the GV is assumed to affect all *m* traits similarly: in this case, the *m* traits should be measured on, or be transformed to the same scale). Finally, **E** is a N × *m* matrix of individual- and trait-specific zero-mean residuals, also referred to as error or disturbance terms. Generally, homoscedasticity of the residuals is assumed, and the *m* × *m* symmetrical background covariance matrix is denoted as *E*[**E**^t^**E**] = **Σ_E_**. That is, **Σ_E_** is the residual variance-covariance matrix between the *m* traits conditional on the *k* predictors, i.e., **E** captures all sources of residual (co)variability. Note that matrix **Σ_E_** is usually not diagonal because, conditional on the *k* predictors, the *m* traits are generally still correlated. Regression-based MATs mainly differ in their treatment of **Σ_E_** (see Box 2). As given in Eq. 2, the multivariate multiple regression model is thus a system of univariate regression equations. By combining them all within one model, specific hypotheses can be tested, and the model can be simplified by introducing constraints in matrices **B** and **E**.

The following regression-based techniques are described in Box 2: Multivariate Analysis of Variance (MANOVA), Generalized Estimating Equations (GEE), and MultiPhen (O’Reilly et al., 2012). Assuming an additive codominant genetic model, MANOVA, GEE models, and Linear Mixed Models (LMM, not included in this review, see Box 2) are specific instances of the model presented in Eq. 2. In contrast, the regression-based MAT MultiPhen is based on reversed ordinal regression with the *m* traits as the predictors and the GV as the dependent variable.

### Combination tests

We define a combination test as any test that combines the p-values or test statistics obtained in *m* univariate analyses to test a multivariate hypothesis. The challenge characterizing combination tests is to optimally handle the correlations between the *m* p-values or *m* test statistics, resulting from the phenotypic correlations between the *m* traits. How the information obtained in univariate tests is combined is described in Box 3 for the following tests: Nyholt-Šidák and Bonferroni corrected p-values (min-P_NS_, min-P_Bonf_; Nyholt, 2004), the Simes test (Simes,1986), its adjusted version TATES (Trait-based Association Test that uses Extended Simes; Van der Sluis et al., 2013), two version of JAMP (Joint genetic Association of Multivariate Phenotypes: JAMP_mult_ and JAMP_min_ (ctg.cncr.nl/software/), the meta-analysis inspired techniques S_Hom_ and S_Het_ (Zhu et al., 2015), and the adjusted Fisher-combination test FC-Pearson (Yan et al., 2016).

We emphasize the following important aspects of these combination tests. First, only 4 of the combination tests truly create, based on the univariate test statistics, a new multivariate test statistics, and, as such, evaluate the *joint* association signal of the *m* traits to the GV (JAMP_mult_, S_Hom_, S_Het_, FC-Pearson). The others essentially constitute various types of corrections for multiple testing. Second, Simes, TATES, min-P_Bonf_, min-P_NS_ and JAMP_min_ do not create a new test statistic, but simply select the smallest of *m* weighted univariate p-values. Due to the weighting (i.e., effectively a correction for multiple testing), the p-values of these combination tests are always larger than the original univariate p-values on which they are based.

These three classes of MATs are conceptually distinguished. Alternatively, all transformation-based and regression-based tests, and some combination tests, can be described from a maximum-likelihood perspective, and within this framework, one could distinguish 1-df and *m*-df tests. Specifically, 1-df tests either reduce all *m* traits to a single new variate (i.e., sum-score analysis, PCA using PC1 only, and factor scores obtained in a single common factor model), or constrain all *m* associations between the GV and the *m*-traits to be equal (S_Hom_, and the 1-df versions of GEE and MANOVA). In all these tests, the association between the GV and the *m* traits is modelled via 1 parameter, which can be tested using a (1-df) likelihood ratio test. In contrast, in *m*-df tests, the associations between the GV and the *m* traits are allowed to vary, and the *m* parameters are subjected to a *m*-df likelihood ratio test, or a closely related (F-) test (standard MANOVA, CPC, and the *m*-df versions of GEE). An alternative classification, based on the underlying mathematical model and the structure of the resulting test statistic, that matches distinction of MATs based on degrees of freedom, is outlined in the Supplemental Information.

Irrespective of their statistical foundation, all MATs need to deal with the fact that the *m* simultaneously modelled traits are often correlated conditional on the tested GV. The way they do so differs: combination tests use either permutation or a correction factor, regression-based tests either treat the *m* traits as predictors, avoiding the issue altogether (MultiPhen), or accommodate the residual trait correlations in a background covariance matrix **Σ_E_** (MANOVA, GEE, LMM), and transformation-based tests explicitly use the covariance between the *m* traits to create the new variate.

## 2. Characteristics of MATS

The classification in transformation-based tests, regression-based tests, and combination tests is based on the statistical properties of the MATs. They differ, however, in various respects that have a bearing on the their performance and applicability. We discuss these differences briefly, and refer to Table 1 for an extensive summary.

### Specific hypothesis tested

While all MATs evaluate the statistical relationship between *m* traits and a GV, they differ with respect to the exact hypothesis that they test. First, MATs can evaluate the omnibus hypothesis that the *joint* association signal of the *m* traits to the GV deviates significantly from 0. This omnibus test can be an *m*-df test, allowing for heterogeneity in the *m* GV-effects regarding sign and size. By assuming the GV-effects to be homogeneous across the *m* traits, the omnibus test reduces to a 1-df test, which can be more powerful if the homogeneity assumption holds approximately. The 1-df tests are obtained through constraining of model parameters (e.g., the regression weights are constrained to be equal), or through the use of transformation-based techniques, in which the *m* traits are reduced to a single new variate under the assumption that this new variate is representative of what the *m* traits have in common. Second, MATs can test the hypothesis that at least one of the *m* traits is significantly associated with the GV. These MATs generally concern combination tests that evaluate the smallest of *m* weighted p-values as obtained in univariate GWA analyses.

### Measurement level of the m traits

The choice of MAT is often largely dictated by the *measurement levels* of the *m* traits. Specifically, if all *m* traits are continuous (to reasonable approximation), PCA, CPC, and MANOVA can be used directly. All MATs suited for continuous data assume the data to be multivariate normally distributed. GEE-based generalized linear modeling can handle continuous or categorical traits, but current standard implementation (e.g., GEE in SPSS or the R library *gee*) cannot handle a mix of different measurement levels. Which measurement levels factor analysis can handle, depends on the software package (e.g., when conducted in MPlus (Muthén & Muthén, 2017) or OpenMx (Neale et al., 2016), factor analysis can in principle handle all measurement levels as well as a mix). The sum score method is applicable to continuous variables, or ordinal variables (including dichotomous) variables (i.e., “burden score”), as long as all *m* aggregated traits are measured on the same scale. If the *m* traits have different measurement levels, combination tests and MultiPhen can be used (but see Guo et al., 2015 on power losses in MultiPhen when traits are non-normally distributed). The strength of combination tests lies in their flexibility to combine results regardless of the traits’ measurement level. For instance, TATES has been shown to work well on a mix of non-uniformly correlating dichotomous, ordinal, and continuous traits (Van der Sluis et al., 2013). The current implementation of the permutation-based combination tests of JAMP is suited for continuous data only, but is in principle amendable to traits with a mix of measurement levels.

### Missingness

In univariate analyses, missing values simply result in a smaller effective sample size N. In a multivariate context, however, partial missingness can occur, i.e, participants having missing values on a subset of the *m* traits. Not all software can handle partial missingness; methods often resort to listwise deletion, basing analyses only on cases with complete data. As in practice the probability of at least 1 of the *m* scores being missing increases with *m*, listwise deletion can result in a substantial reduction of sample size and consequently a considerable reduction in statistical power. Alternatively, however, one can use packages like OpenMx (Neale et al., 2016) that use Full Information Maximum Likelihood (FIML, i.e., all available data are used) to specify a wide variety of multivariate models (including MANOVA, PCA, and factor analysis) while accommodating the missingness. This can, however, come with a prohibitive computational burden in the GWA settings.

If one weights the *m* trait scores appropriately, sum scores can still be used if the data show partial missingness: e.g., each individual sum score may be divided by the number of observed trait scores. As this may result in heteroskedastic variance, weighted sum scores are generally used in combination with a cut off criterion (e.g., no more than 20% of the *m* scores can be missing), which also ensures approximate conceptual comparability between scores over subjects with different numbers of observed scores.

The essentially univariate nature of the input of combination tests guarantees their ability to handle missingness. However, if sample sizes differ greatly between the *m* traits, a (sample size) weighted procedure (like S_Hom_ and S_Het_ offer) is desirable.

Generally, partial missingness lowers the power to detect GV, especially if the traits with a relative large percentage of missingness are the traits with the strongest genetic association. Additionally, in using methods that can accommodate the missingness, one should realize that the multivariate association signal may be primarily driven by the traits with the lowest percentage of missingness.

Imputation of the missing scores can be a convenient way to handle missing data, as replacement of the missing values with imputed ones facilitates the use of all MATs. Multivariate imputation, i.e., dealing with imputation of missing values in multiple variables at once, can be done in many ways, but comes with its own challenges and can yield biased results (see e.g. Nakai & Ke, 2011; van Buuren & Groothuis-Oudshoorn, 2011).

### Relateds

GWA data sets may include data collected in families (e.g., trios of parents and one affected off-spring, data of twins and their family members). In univariate analyses, inclusion of family members can be useful to differentiate “between” from “within” family associations, the latter being free of any effects of population stratification (Fulker et al., 1999). Also, including all available data, even data of genetically similar monozygotic twin pairs, can be beneficial in terms of power to detect GV-effects (e.g., Minica et al., 2014). However, if data include family members, the data clustering induced by the relatedness must be accommodated statistically to avoid inflated Type I error rates. In the univariate setting, multiple linear mixed model approaches exist (see Eu-ahsunthornwattana et al., 2014 for comparisons). When data only include a few relateds, one can chose to “correct for” the familial relatedness rather than explicitly model it. For instance, PLINK (Purcell et al., 2007) offers the option to correct for relatedness in the data by running GEE, which involves a correction of standard errors^1^. In principle, these univariate procedures can be used in the context of transformation-based techniques (i.e., correcting the univariate analyses of the new variate), and in the context of combination tests, in which case the corrected model parameters of the *m* univariate GWA analyses are used as input for the combination tests (to our knowledge, only the performance of the combination test TATES has been studied in the context relatedness; Vroom et al., 2015). Combination tests using permutation, like JAMP, need to permute the data not on an individual level but on the family-level to retain the familial relatedness in the data. This is complicated if the families in the data set do not all have the same size and composition.

In their standard form, MANOVA and MultiPhen cannot be used on data including relateds. Theoretically, in case of familial clustering, multivariate multilevel modelling can be used instead of MANOVA (Pituch and Stevens, 2016), and Structural Equation Modelling can be used instead of MultiPhen, treating the *m* traits as exogeneous variables. These approaches are, however, computationally intensive.

As standard GEE software can handle only one source of clustering at the time, it can handle either familial relatedness in a univariate setting, or multivariate data in a sample of genetically unrelated individuals, but not both. In principle, LMM (Box 2) can handle multiple sources of clustering or correlation.

### Computational feasibility

Given imputation of genetic variants, current GWA studies may include tens of millions of SNPs. Cluster computers offer large computation capacity, but computation burden is an important consideration in the choice of MAT. In theory, any of the MATs discussed here can be applied using standard software. However, in practice, the use of dedicated software like PLINK (Chang et al, 2015, Purcell et al, 2007) considerably facilitates running such vast amounts of statistical tests on files containing multiple terabytes of data. From a computational feasibility perspective, MATs that rely on univariate analyses (i.e., transformation-based tests and combination tests) or MATs that are built-in in dedicated software (Canonical Correlation Analysis, i.e., MANOVA (see Box 1) as part of PLINK) may be preferred over tests like GEE, MultiPhen, S_Hom_ and S_Het_, or permutation-based tests like JAMP_mult_ and JAMP_min_. Due to their increased computational intensity, these latter options are particularly attractive if they indeed come with clear advantages, like substantial gains in power.

## 3. Type I error rates of MATs

A correct Type I error rate is a primary requirement of any statistical test. We studied the Type I error rates of 17 MATs, excluding the JAMP-methods as the correctness of their Type I error rates is guaranteed by their reliance on permutation. The 17 MATs were studied in 20 scenarios that are outlined in Table 2 (see Supplemental Information for simulation details). The 20 scenarios varied with respect to the number of included variables (*m*=4 or *m*=16), the strength of the correlations between the traits, and the correlational structure, i.e., uniformly correlated traits (i.e., 1-factor model with compound symmetry), or two clusters of more or less strongly correlated traits (i.e., 2-factor model). All simulated traits were standard normally distributed. For each scenario, we ran N_sim_=1,000,000 replications, allowing us to reliably evaluate Type I error rates at α-levels of .05, .01, and .001. All Type I results are available in Tables S7-S9.

**Table 2.**
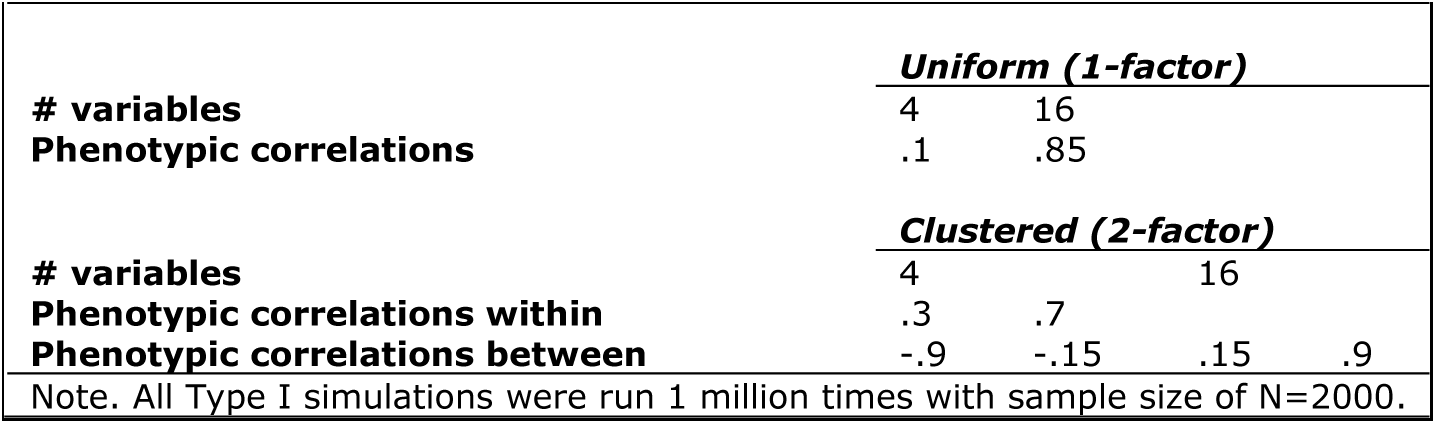
Overview Type I simulation settings.

|  | *Uniform (1-factor)* |  |  |  |
| --- | --- | --- | --- | --- |
| # variables | 4 | 16 |  |  |
| Phenotypic correlations | .1 | .85 |  |  |
|  | *Clustered (2-factor)* |  |  |  |
| # variables | 4 |  | 16 |  |
| Phenotypic correlations within | .3 | .7 |  |  |
| Phenotypic correlations between | -.9 | -.15 | .15 | .9 |

Note. All Type I simulations were run 1 million times with sample size of N=2000.

We note that the large number of replications provides high statistical power to detect small deviations from the expected Type 1 error rate (α), especially for the larger α values. For instance, with 1 million replications, the 99% confidence interval (CI_99_) for α=.05 is very narrow: .04944-.05056 (see Table S6 for the CI_99_ for all α-levels). As a result, merely considering which MATs show Type I errors outside the CI_99_ paints a gloomy picture (Figure S2a). Type I error rates of MANOVA, S_Hom_, and all transformation-based (i.e., essentially univariate) MATs are virtually always correct. However, when considered across all 20 scenarios and all three levels of α (.05, .01, .001, i.e., 60 scenarios in total), all other MATs showed Type I error rates outside the CI_99_, with overall percentages ranging from 22% (CPC) to 92% (FC Pearson) and 100% (GEE_uns_*m*_).

Figure 1 shows the Type I error rates of the 17 MATs given α=.05 for 4 or 16 variables, split for scenarios with mostly low or mostly high trait correlations (see Table S2). As many of these deviations outside the CI_99_ were (very) small (Tables S7-S9), we also looked beyond the CI_99_ by summing the deviations from the expected α across all scenarios, allowing us to determine which factors caused the largest deviations (Figure S2b). Overall, the largest deviations are observed for TATES, min-P_NS_, Simes, FC-Pearson, GEE_uns_*m*_, and min-P_Bonf_. Interestingly, combination tests show mainly deviations from the expected when the *m* traits are highly correlated, while the number of traits *m* mainly drives the deviations in most other method. Taking the direction of the deviations into account, we see that CPC, Simes and min-P_Bonf_ are always conservative, while S_Het_, Tates and min-P_NS_ are conservative when applied to many (highly correlated) traits, and liberal otherwise. All other methods that do show deviations from the expected, always show inflation, with Type I error rates of GEE_uns_*m*_ and FC-Pearson especially being inflated when *m* is large, irrespective of the correlations between the phenotypes.

**Figure 1:**
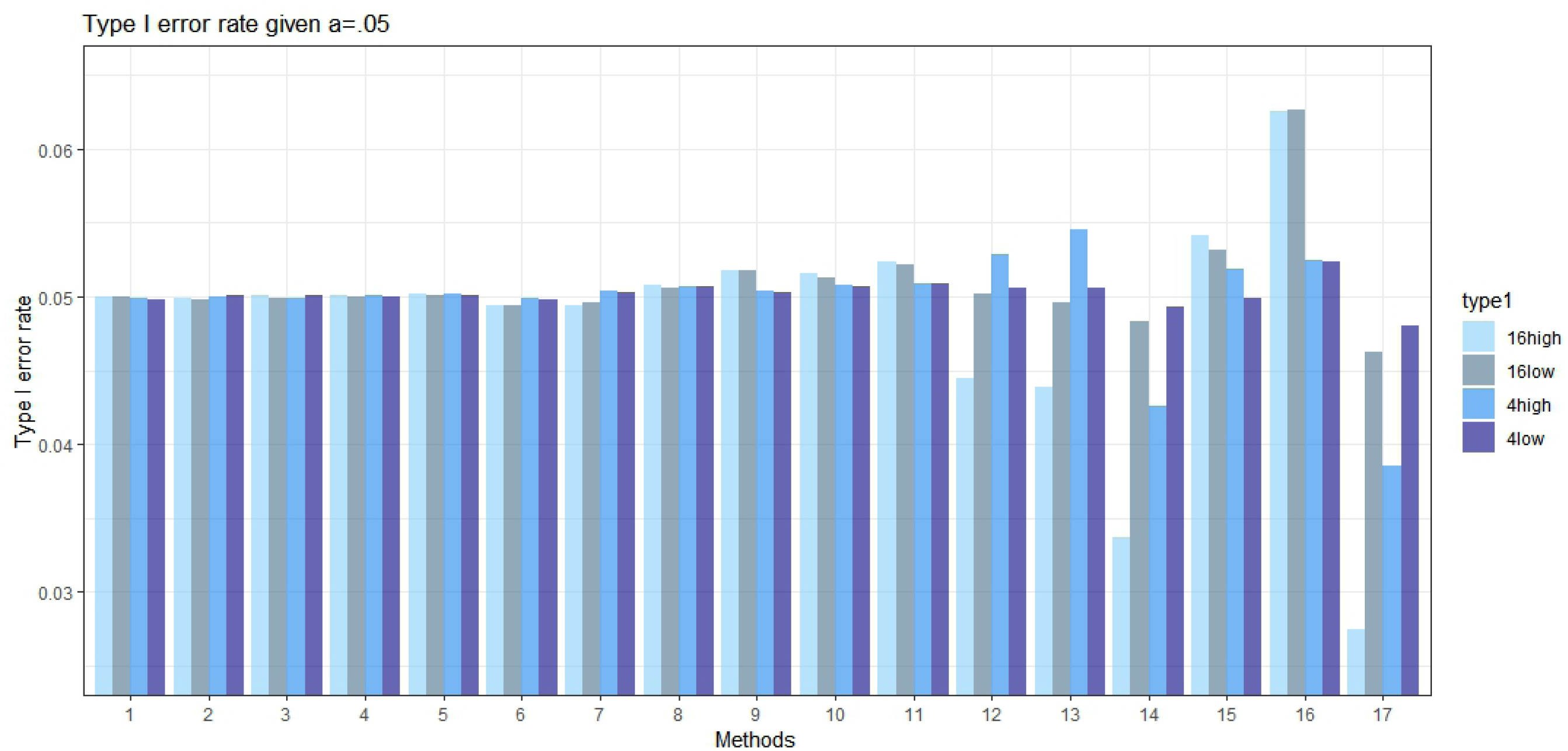
>Type I error. Type I error rates for 17 MATs given Nvar=4 or Nvar=16, plotted separately for scenarios with mostly low and scenarios with mostly high correlations (see Supplemental Table S2). Methods are numbered: 1=MANOVA, 2=factor score, 3=PCA, 4=sum score, 5=SHom, 6=CPC, 7=SHet, 8=GEEex-1, 9=MultiPhen, 10=MANOVA 1df, 11=GEEun-1, 12=TATES, 13=min-PNS, 14=Simes, 15=FCPearson, 16=GEEun-m, 17=min-PBonf. See Supplemental Tables for Type I error rates given α=.01 and α=.001. Note: the two JAMP-methods were excluded from the Type I error rate study as the correctness of their Type I error rates is guaranteed by their reliance on permutation.

Summarizing, due to the strong power to detect deviations from the expected, many methods showed Type I error rates outside the CI_99_. When considering the magnitude of the deviations, especially application of Simes, min-P_Bonf_, FC-Pearson, and *m*-df versions of GEE warrant careful consideration, although even here the actual deviations are often quite small (Tables S7-S9).

## 4. Power of MATs

The statistical power of a test is the probability that the null-hypothesis of no association is correctly rejected when the GV is indeed statistically associated with the trait(s). In de context of GWA studies, GV-effects are expected to be small, so in selecting a MAT for one’s analyses, power is an important consideration.

We studied the power of 19 MATs in 15 scenarios covering 270 settings of the true genotype-phenotype model, which are summarized in Tables 3 and 4 (see Supplemental Information for simulation details). The scenarios varied with respect to the number of traits (*m*=4, 8, or 16, all standard normally distributed), the correlational structure (i.e., uniformly correlated or clustered, corresponding to 1- or 2-factor models), the strength and sign of the correlations between the *m* traits, the number of traits affected by the GV (1, half, or all *m*), and the presence or absence of opposite effects (i.e., GV affecting multiple traits but in opposite direction). For each setting, we ran 1,000 simulations with a GV explaining .1, .2 or .5% of the variance in each affected trait, and a sample size of N=2000.

**Table 3.**
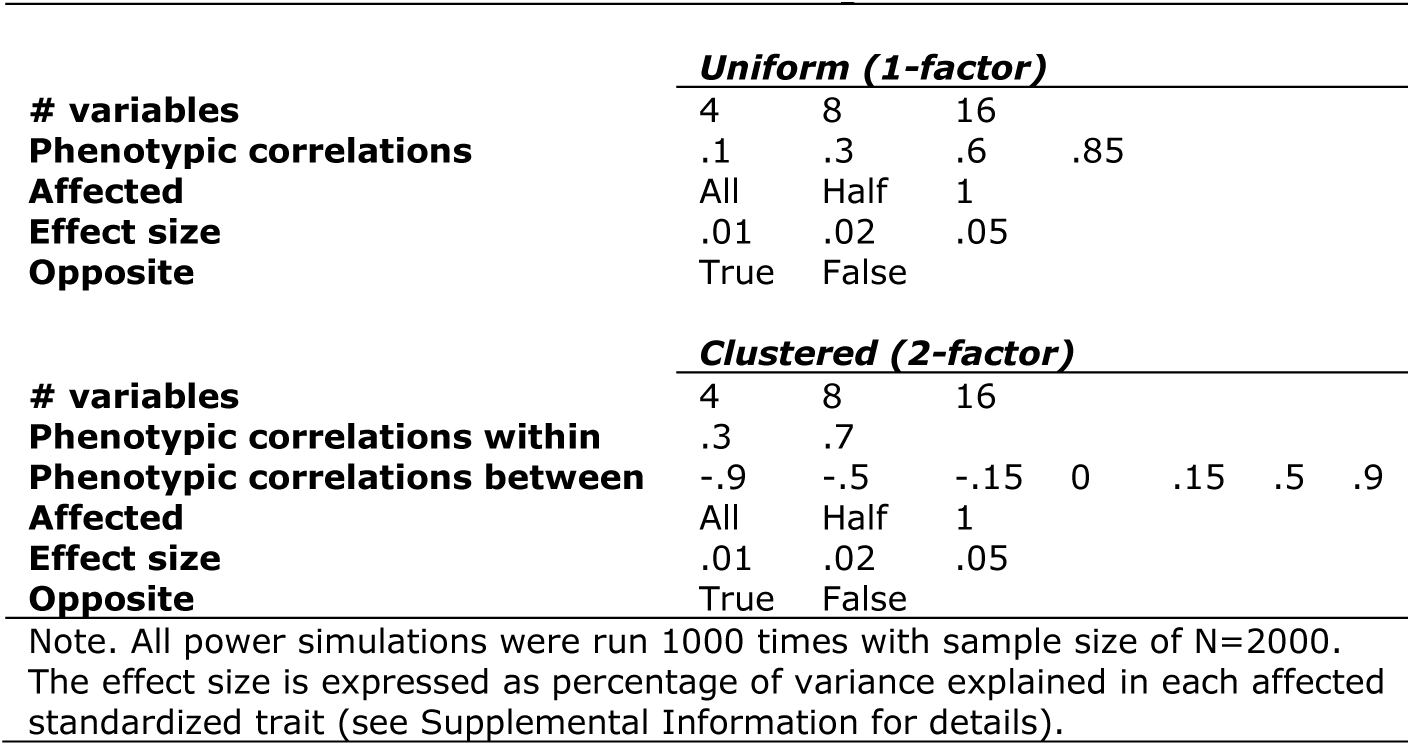
Overview Power simulation settings.

**Table 4.**
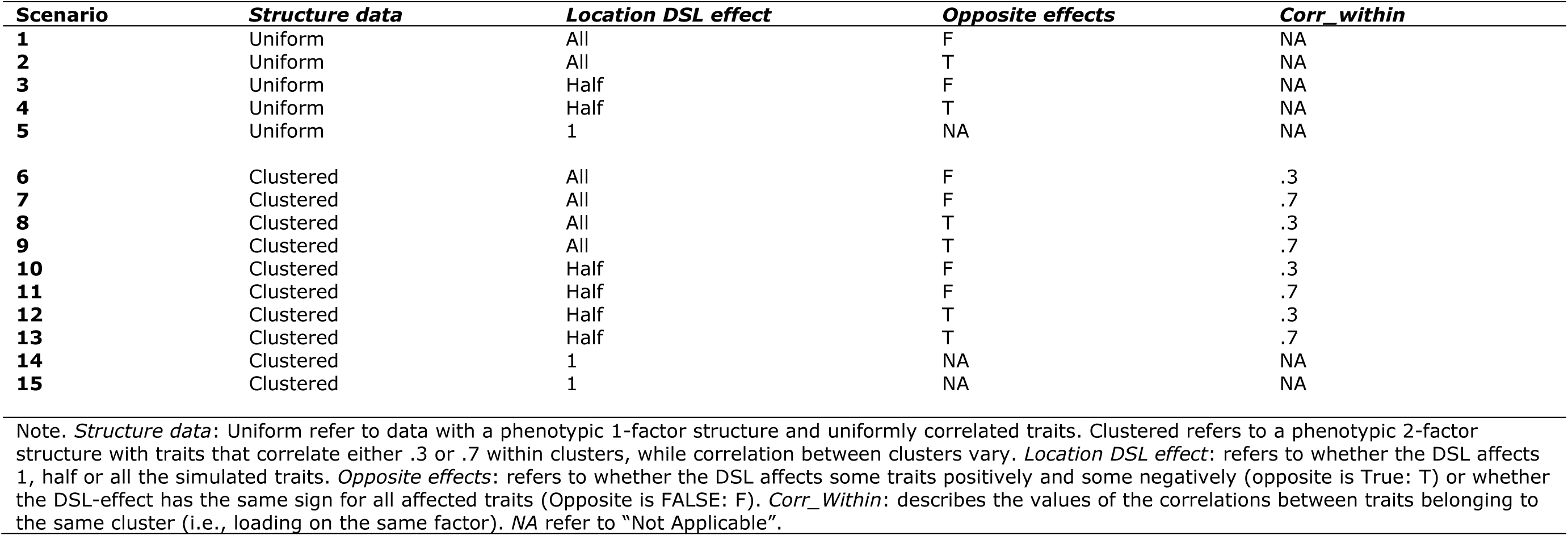
Overview 15 main power simulation scenarios.

The full results of the power simulations are available in Table S10-S12. Below, we discuss the power results for a GV explaining .1% of the variance (Table S10), and emphasize that these main finding hold for GV of different effect sizes (Tables S11-S12). We excluded the 2 MATs with highly inflated Type I error rates (GEE_uns_*m*_, and FC-Pearson) from discussion as their power estimates can be biased upwards due to the inflated Type I error rates (but see Tables S10-S12 for all power results of these test). We did include the two conservative MATs (Simes, min-P_Bonf_) in our discussion, as their deflated Type I error rates will result in under- rather than overestimation of power which we can interpret as a lower bound estimate.

Figure 2 depicts the power of these 17 MATs in all 15 scenarios for 4 and 16 variables. We note that the power of MATs can be compared within, but not always directly between, scenarios as the total contribution of the GV to the *m* traits can differ across scenarios as a function of the correlations between the *m* traits.

**Figure 2:**
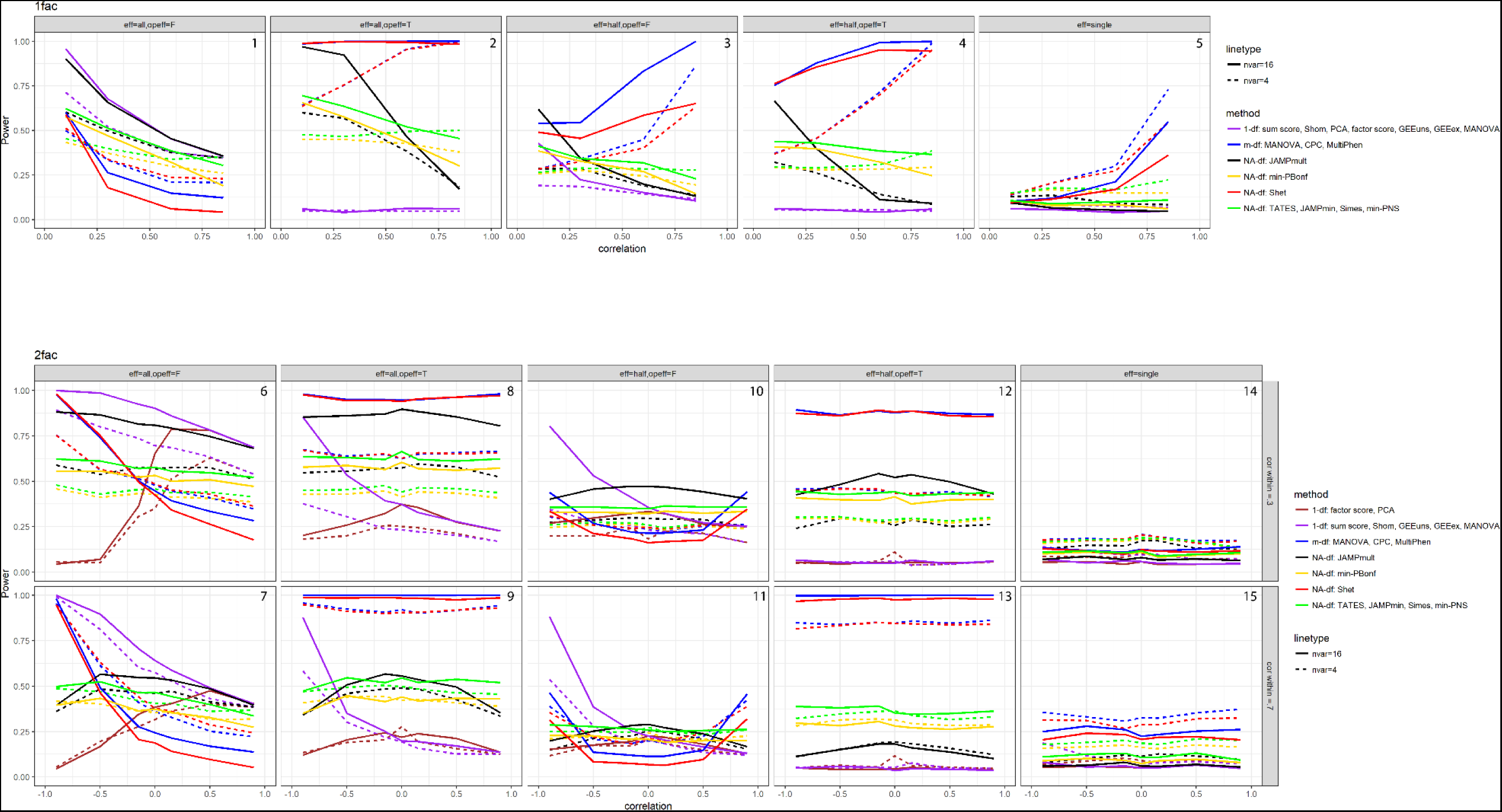
Power. Panels **a** and **b** show the power to detect a GV that explains .1% of the variance (see Supplemental Information) as a function of the number of traits (4 or 16; see Table S10 for results for 8 traits) and the correlations among the traits. Power curves are shown for 17 MATs in the 15 scenarios outlined in Table 4.

### Univariate versus multivariate

When testing the association of a GV to *m* traits, one could simply do *m* univariate analyses and correct the *m* resulting p-values for multiple testing. We consider the power results of the combination test min-P_Bonf_ an approximation of this approach (although min-P_Bonf_ subsequently selects the smallest Bonferroni corrected p-value). The power results in Figure 2 reveals that when all or half of the *m* traits are affected by the GV (scenarios 1-4, 6-13), MATs are very often (but not always!) more powerful than a for multiple testing corrected univariate analysis. MATs even often outperform univariate analyses when only 1 of the *m* trait is affected by the GV, especially when the trait correlations are generally high. Taken over all scenarios, it is safe to conclude that multivariate approaches towards identification of GV are generally worth pursuing.

### Equivalence of MATs

So far, we classified MATs based on their underlying statistical approach, the descriptions in Boxes 1-3 outlining their differences. The power simulations, however, demonstrate that there are 3 groups of MATs that function very similarly, i.e., have very similar power across all or most of the scenarios (see Supplemental Information for detailed comparisons). First, the combination tests min-P_NS_, Simes, TATES, and JAMPmin demonstrate very similar power throughout all 15 scenarios, with min-P_Bonf_ showing a very similar yet consistently lower power profile. Second, the *m*-df tests MANOVA, CPC, and MultiPhen perform very similarly (and very similar to the *m*-df variants of GEE), with S_Het_ generally does equally well or slightly worse. Third, in the context of uniformly correlated traits (scenarios 1-5), tests that can generally be referred to as 1-df tests group together, i.e., the transformation-based techniques sum-score, PCA, and factor scores, and the 1-df variants of the regression-based tests GEE (exchangeable and unstructured) and MANOVA. However, in the context of clustered traits (scenarios 6-15), PCA and the factor scores perform much worse than the other 1-df tests when the clusters correlate negatively. Interestingly, the combination test JAMP_mult_ follows its own trend (which is very similar to that of the FC-Pearson test).

### Relative insensitivity to the true genotype-phenotype model

The true genotype-phenotype model provides the multivariate context in which one tests the associations between the *m* traits and the GV. Our power simulations show that some MATs are relatively insensitive to this context, i.e., their power varies much less across the different scenarios compared to other MATs. These relatively insensitive MATs all concern combination tests that are based on selection of the minimum weighted p-value: min-P_Bonf_, min-P_NS_, Simes, TATES, and JAMP_pmin_. Mainly in the context of many uniformly correlated traits and a *pleiotropic* variant affecting all *m* traits (scenarios 1-2), do these methods demonstrate noticeable variation in power, i.e., their power to detect the GV decreases with increasing correlations between the *m* traits, irrespective of the presence of opposite effects. In all other scenarios, the power curves for these methods are rather flat, illustrating their relative insensitivity.

This relative insensitivity to the true genotype-phenotype model can be advantageous: there are several settings in which these MATs generally outperform *m*-df tests and S_Het_ (e.g., scenarios 1, 6, 7, 10, 11), factor scores and PCA (e.g., scenarios 6-9, 12 and 13), JAMP_mult_ (3-5,13-15), and sum scores, S_Hom_ and 1-df regression-based tests (e.g., 2-5,8,9,12 and 13). However, some MATs actually benefit from specific characteristics of the true genotype-phenotype model, such as the presence of unaffected or oppositely affected variables in the analysis (see below). Under these circumstances, these relative insensitive MATs are, sometimes substantially, outperformed. Because of their relative insensitivity, we exclude these MATs from further discussion.

### Clustered versus uniformly correlated traits

When the *m* traits are uniformly correlated, all transformation-based techniques have very similar power (scenarios 1-5). In this context, the power of transformation-based techniques increases with decreasing correlation among the *m* traits. Specifically, the variance of the new variates, summarizing the communality between the traits, is larger when the *m* traits correlate more strongly and the contribution of the GV to that common variance is in that case relatively small. That is, the *signal-to-noise ratio* is more optimal when the covariance between the traits conditional on the GV is low (see Supplemental Information for an elaborate discussion).

In the context of clustered correlated traits, however, PCA and factor scores perform differently from the other transformation-based tests when the correlation between clusters of positively correlated traits is negative (scenarios 6-11). In that case, the first PC from PCA and the factor scores from a 1-factor model will only summarize 1 of the two clusters well, while they do not capture information from the other cluster. Interestingly, in the calculation of sum scores, the presence of negatively correlated variables can actually have a beneficial effect on the detection of GV-effects (scenarios 6-11): the negative covariances between pairs of traits reduce the total variance of the sum, which in turn improves the signal-to-noise ratio (see Supplemental Information).

When the GV affects only half or 1 of the traits, the *m*-df tests MANOVA, MultiPhen and CPC perform better when the *m* traits are uniformly correlated than when they are clustered (scenarios 3 and 5 versus 10-11 and 14-15), but when the GV affects all *m* traits or conveys opposite effects (scenarios 1,2,4 versus 6-7,8-9,12-13), the power of these tests does not seem to suffer much from the clustering in the data.

In the context of uniformly correlated traits (scenarios 1-5), the power of JAMP_mult_ is clearly a function of the trait correlations, with lower trait correlations resulting in higher power. Similar results are observed for the clustered scenarios, if one compares the power in the scenarios with within-cluster correlations of .3 (scenarios 6,8,10,12 and 14) to those with within-cluster correlations of .7 (scenarios 7,9,11,13 and 15: always lower).

### Pleiotropic versus local variants

In evaluating GV-effects in a multivariate context, it is desirable to distinguish between the detection of pleiotropic or *global genetic variants* (i.e., variants that affect all or multiple of the *m* traits in the analysis) and *local genetic variants* (i.e., variants that effect only 1 or a few of the *m* traits in the analysis). As we defined a MAT as any test that formalizes the statistical association between a GV and a set of *m* traits that are measured in the same individual, one may argue that MATs should be assessed based on their power to detect global variants. Conducting multivariate analyses may then not only be lucrative with respect to power, but can also aid theoretical development and biological understanding by revealing shared underlying biology. However, a one-sided focus on global variants neglects the importance of identifying local variants, which may be a source of genetic heterogeneity. Identification of genetically homogeneous subsets of traits within the full set of *m* traits acknowledges the contribution of more local variants and may be biologically informative (e.g., Nagel et al., 2018).

In the context of uniformly correlated traits, the (transformation-based) 1-df tests work best for the identification of global variants that affect all phenotypes in the same direction (scenario 1), as these contribute most to the variance of the new variate. Here, the power to detect global GVs decreases as the conditional correlations between the *m* traits increase (i.e., the *signal-to-noise ratio* decreases). Yet, GVs that affect only half or 1 of the *m* traits (scenarios 3, 5) can hardly be detected through these 1-df tests: such GVs will generally contribute little to the variance of the new variate and will therefore be (very) difficult to identify using transformation-based approaches. When traits show clustering, we see a clear difference between sum scores and other 1-df MATs, which do well in detection global variants (scenarios 6,7), and PCA and factor scores, which do poorly. Clearly, the first PC and factor scores based on a 1-factor model do not capture the clustered nature of the data well. Interestingly, in a clustered context, 1-df tests do best in detecting GV affecting only half of the *m* traits (scenarios 10,11), especially when the unaffected traits correlate negatively to the affected traits: in that case, the negatively correlations lower the variance of the new variate and as such improve the signal-to-noise ratio. Yet, truly local variants go undetected when transformation-based or 1-df MATs are used.

Conceptually, MATs that evaluate the joint association signal of the *m* traits through *m*-df omnibus tests truly test for global variants, i.e., Cross Phenotype (CP) associations, i.e., whether a genetic variant is associated with more than one trait (i.e., pleiotropic, see Solovieff et al., 2013). Counter intuitively, however, our simulations demonstrate that in the context of both uniformly correlated and clustered traits (scenarios 1,6,7), those *m*-df MATs do not have the best power to detect global variants, and (like for all MATs) their power suffers especially when the *m* traits correlate substantially (Minica et al., 2010; Medland & Neale, 2010). When traits correlate uniformly, these *m*-df MATs do have the best power to detect local GVs (scenario 5) and GVs that affect only half of the *m* traits (scenarios 3). In case of clustered variables, the presence of negatively correlated variables can boost the power to detect global GVs (scenarios 6,7), but their power to detect GVs that affect only half (scenarios 10,11) or 1 (scenarios 14,15) of the *m* traits is generally very low, although still superior to that of other MATs.

JAMP_mult_ is quite good at picking up global GVs, especially when the trait correlations are low (scenarios 1-4,6). In the context of uniformly correlated traits, JAMP_mult_ has noticeably less power than the *m*-df tests to pick up GV that affect only 1 or half of the *m* traits, especially with increasing correlations between the *m* traits. In clustered settings, JAMP_mult_ can perform slightly better than *m*-df tests when GV affect only half of *m* traits (e.g., scenarios 10,11).

### Presence of unassociated traits

In psychology and clinical research, it is common to observe mean group differences in some but not all variables of a set of *m* moderately/highly correlated traits. For instance, Van der Sluis et al (2008) observed significant gender differences in the means of 3 out of 12 substantially positively correlated cognitive subtests of the WISC-R (Carroll, 1993). Similarly, gender differences in endorsement rates are often observed in some but not all of positively correlated depression symptoms (see e.g. Lux & Kendler, 2010). In genetic research, where GV-effects are generally small, it is likely that a GV affects correlated traits differently. For instance, in a set of 12 phenotypically correlated neuroticism items (.17-.54), Nagel et al (2018) identified many item-specific genome-wide significant genetic regions (see their Supplementary Data 2). As the exact GV-trait relationship is generally unknown, it is important to consider the effect of the presence of unassociated traits in the set of *m* traits on the power of MATs.

To study the effect of the presence of unaffected traits on the power to detect as GV of interest, we compare the power results of scenario 5 for 4, 8 and 16 variables, i.e., the power to detect a local GV-effect in the presence of 3, 7, or 15 unaffected variables, respectively (Table S10). In this context, the power to detect the GV is low for all methods, except the *m*-df techniques MANOVA, MultiPhen and CPC, and S_Het_, which do have some power if the trait correlations are substantial (i.e., .5 or higher). For all MATs, the power to detect that local GV deteriorates when more unaffected uniformly correlated traits are added to the analysis.

Interestingly, the *m*-df tests MANOVA, MultiPhen and CPC, and S_Het_ have lower power to detect a GV that affects all *m* traits (scenario 1) than to detect a GV affecting half of the *m* traits (scenario 3), even though the total amount of signal is lower in the latter scenario. Specifically, the presence of unaffected traits can boost the power to detect GV effects considerably, but only if they are substantially correlated to the affected traits in the analysis. In the Supplemental Information, we show graphically for *m*=2 (inspired on Cole et al., 1994) how a GV that affects trait Y1 but not trait Y2 can aid discrimination between genotype groups (and thus detection of the GV).

### Opposite effects

GV with opposite effects, in which an allele increases the value of/risk to one trait, while decreasing the value of/risk to another, are not uncommon (Solovieff et al., 2013). For instance, Sitora et al (2009) demonstrated such opposite effects in autoimmune diseases. Given the existence of GVs with opposite effects, it is important to determine which MATs can detect them.

Our simulations show that the power of all 1-df MATs (both reduction and regression-based techniques, and S_Hom_) suffers seriously from the presence of opposite effects. The transformation-based tests all rely on the variance that is shared between the *m* traits, i.e., their communality. While concordant effects contribute to this communality, opposite effects do not and cancel out. Consequently, the opposite GV-effects are poorly represented in the new variate (depending on the ratio concordant-to-opposite effects), thus resulting in decreased power to detect them.

Under the assumptions that the GV-effects are concordant across all *m* traits, 1-df MATs constrain them to be equal and then test whether this single parameters deviates significantly from 0. When the assumption holds, this reduced model has increased power to detect the GV compared to univariate procedures (e.g., scenarios 1,6,7). However, if the GV-effects are opposite in reality, constraining them to be identical will cancel individual effects out, thus drastically reducing the power of 1-df MATs (e.g., scenarios 2,4,12,13). Interestingly, when clusters of traits correlate negatively (e.g., scenarios 8,9), the GV-effects can contribute to the communality if the difference in sign of the GV-effect is in concordance with the difference in sign of the correlations, in which case GV with opposite effects can be picked up by these methods.

In contrast, JAMP_mult_ handles opposite effects much better than transformation-based and 1-df tests, while the *m*-df MATs MANOVA, MultiPhen, and CPC, and S_Het_ actually seem to benefit from the presence of opposite effects (scenarios 2,4,8,9,12,13). That is, the power to identify opposite-effect GVs that affect all or half of the *m* traits is actually higher than the power to detect a GV that has concordant effects on half or all of the *m* traits (pairwise compare scenarios 1 to 2, 3 to 4, 6 to 8, 7 to 9, 10 to 12, 11 to 13). As *m*-df tests evaluate the *m* association parameters individually, the effects do not cancel each other out. Cole et al (1994) already showed that for MANOVA, the critical consideration is not simply the sign of the GV-effects, but the sign of the correlation between the traits as well. In the Supplemental Information, we show graphically for *m*=2 (inspired on Cole et al., 1994) how a GV that increases the mean of trait Y1 while decreasing the mean of trait Y2 can aid discrimination between genotype groups (and thus detection of the GV) if these traits are positively correlated.

### Number of traits

In planning multivariate analyses, one important question is whether the power to detect the GV depends on the number of traits. Our simulations show that when the GV affects only 1 of the *m* traits, the power is generally slightly better if *m* is smaller (scenario 5,14,15; Figure 2). For all other scenarios (i.e., scenarios concerning GVs that affect half or all of the *m* traits), including more traits is generally beneficial for the power of all MATs, except for the *m*-df tests. For the *m*-df tests, including more traits is only beneficial when the GV transmits opposite effects (scenarios 2,4,8,9,12,13). Yet, when a GV affects all of the *m* traits similarly (scenarios 1,6,7), or only 1 of the *m* traits (scenarios 5,14,15), then these *m*-df tests have better power when *m* is small because in that case the number of degrees of freedom is smaller.

## 5. Discussion

Researchers often employ MATs with the aim to discover pleiotropic GVs, i.e., GVs that are statistically associated to multiple traits, which possibly points towards a shared biological substrate (Solovieff et al., 2013). The general finding of our simulations that the power to detect such global variants decreases for all MATs as the phenotypic correlations between the traits increase (e.g. Minica et al., 2010, Medland & Neale, 2010; as would be expected with increasing genetic relatedness), demonstrates that currently available MATs are actually not optimised to identify true pleiotropic GVs (see also Porter et al., 2017).

The considerable variation in power displayed by MATs across multiple scenarios demonstrates that the choice of MAT is no trivial matter. The optimal choice is determined by multiple factors that define the true genotype-phenotype model, such as the strength and sign of the correlations between the traits, sign and generality of the GV-effect, and the presence of unaffected traits. Many of these factors are unknown prior to analysis, which hampers the formulation of globally applicable recommendations. As Zhou & Stephens (2014) noted “…in a GWAS setting no single test will be the most powerful to detect the many different types of genetic effects that could occur. Indeed, it is possible to manufacture simulations so that any given test is most powerful. Thus different multivariate and univariate tests should be viewed as complementary to one another, rather than competing.” Consequently, identifying the circumstances in which specific MATs perform strongly or poorly, and indicating which (classes of) MATs are most versatile, is the best we can do for now. Overall, the *m*-df MATs outperform both transformation-based tests and combination tests in 10 out of the 15 scenarios (2-5, 8,9,12-15) represented in our study. That is, the *m*-df MATs are better at identifying GVs that convey opposite effects or GV that affect only a subset of the modelled traits, but are often outperformed when GV are truly pleiotropic (scenarios 1,6-7).

As previously pointed out concerning MANOVA (Cole et al., 1994), the power of *m*-df MATs can, somewhat counter-intuitively, improve from the inclusion of traits that are unassociated to the GV, if these are correlated with the affected traits. In the context of experimental studies, this knowledge can be put to use given prior or theoretical knowledge of which traits are expected to be affected or unaffected by a given manipulation. In the context of GWAS, however, such theory to guide in- or exclusion of traits is usually lacking (see Supplemental Information for a short discussion on the (dis)advantages of increasing the number of traits *m*).

In our simulations, we considered only additive codominant GVs and normally distributed continuous traits. These choices fit the (distributional) assumptions underlying most MATs. We note that Type I error rates of various techniques (e.g., MANOVA, univariate regression) may not be correct when standard assumptions are violated (e.g., severely non-normal or non-continuous data, see e.g. O’Reilly et al, 2012, Yang et al., 2016, Gasperik, 2010), and that some MATs may have better power to identify non-additive GVs than others. Yet for a selection of MATs, Porter & O’Reilly (2017) showed that for those methods amendable to dichotomous case-control data, the pattern of results was remarkably similar to that obtained using continuous data.

In the current review, we focused only on frequentist-based MATs that do not rely greatly on permutation or bootstrapping. MATs based on Bayesian modeling do, however exist (e.g. multivariate version of SNPtest (Marchini et al., 2007) and BIMBAM (Stephens, 2013)) or bootstrapping (e.g., PCHAT, Klei et al, 2008), and we refer to Galesloot et al (2012) and Porter et al (2017) for power simulations including these MATs. Similarly, we focused on MATs that formalize the statistical association between a GV and a set of *m* traits that are all measured on the same individual. Recently, multiple methods were developed that allow estimation of the genetic covariance between traits using genome-wide association signal (e.g., GCTA (Yang et al., 2011), BOLT-REML (Loh et al., 2015), LD Score Regression (Bulik-Sullivan et al., 2015)), alongside multivariate methods like Multi-Trait Analysis of GWAS (MTAG: Turley et al., 2018) and genomic SEM (Grotzinger et al., in press), which use this genetic covariance among traits to boost the statistical power to detect GVs for (sets of) target traits. As these techniques are not primarily SNP-level multivariate tests of traits measured on the same individual (although genomic SEM can be used as such), they were not included in this review.

Summarizing, we presented a classification on MATs based on both their underlying statistical approach and the associated degrees of freedom, alongside a summary of their main characteristics. We showed that MATS vary considerably in their power to detect associated GVs, that under many circumstances, MATs are often more powerful than multiple testing corrected univariate analyses even when only 1 of the *m* traits is affected by the GV, and that in many scenarios *m*-df MATs are the most powerful. We also demonstrated for all current MATs, the power to identify truly pleiotropic GVs decreases with increasing trait-correlations, i.e., particularly when pleiotropy is expected. With increasing availability of multivariate information from large publicly accessible biobanks (e.g., UK Biobank, 23andMe, deCODE), and knowing that pleiotropy is wide-spread both within and between trait domains (Watanabe et al., in revision), we believe that development of new MATs that focus specifically on detection of pleiotropic GVs is crucial. Through sharing of flexible simulation scripts, we facilitate a standard framework for comparing Type I error rate and power of new MATs to that of existing ones.

## Acknowledgement

This work was funded by The Netherlands Organization for Scientific Research (NWO MagW VIDI 452-12-014, NWO VICI 435-14-005 and 645-000-003). Analyses were carried out on the Genetic Cluster Computer, which is financed by the Netherlands Scientific Organization (NWO: 480-05-003), by the VU University, Amsterdam, the Netherlands, and by the Dutch Brain Foundation, and is hosted by the Dutch National Computing and Networking Services SurfSARA.

## Footnotes

1 Specifically, the working correlation matrix is by default set to “independent” in PLINK (i.e., the family scores are assumed independent conditional on the GV under study) to minimize computational intensity. GEE’s standard sandwich correction then corrects the standard errors of all estimated parameters for model misspecification induced by ignoring relatedness. This procedure works well in terms of Type I error rates, but Minica et al (2015, see also Vroom et al., 2016) showed that considerable statistical power can be gained if the working correlation matrix is set to unstructured, although this is computationally more demanding.

2 Note that the JAMP software also calculates an empirical p-value that controls for the family wise error due to testing multiple SNPs. This family-wise corrected p-value tends to be less conservative than the Bonferroni corrected p-value, as it properly takes into account the correlational structure of the genomic data. This family-wise corrected p-value was not used in the current study.

