## Supplemental Information for "The more the merrier? Multivariate approaches to genome-wide association analysis"

#### Table of Contents

| | | |
|---|---|---|
| | | **Page** |
| **S1 – Overall aim** | | **3** |
| **S2 –Selection of MATs for simulations** | | **3** |
| **S3 – Alternative classification** | | **4** |
| **S4 – Simulations** | | **5** |
| **S5 – Type I error rates** | | **12** |
| **S6 – Power results** | | **15** |
| **S7 – Equivalence of GEE *m*-df tests** | | **16** |
| **S8 – The variance of sum scores** | | **18** |
| **S9 – MANOVA** | | **20** |
| **S10 – Adding variables: good or bad idea?** | | **22** |

### S1. Overall aim

The aim of the current review was to investigate and compare the Type I error rate and power of various multivariate association tests (MATs) that are used in genome-wide association (GWA) settings. We define a MAT as any test that formalizes the statistical association between a genetic variant (GV) and a set of *m* traits that are measured in the same individuals. Methods that were specifically developed to conduct multivariate analyses based on univariate GWA summary statistics from potentially different cohorts (e.g., MTAG (Turley et al., 2018), genomic SEM (Grotzinger et al., in press)) are thus not included in the current review.

### S2. Selection of MATs for simulations

The MATs that we included in the current review are described in Boxes 1-3 of the main manuscript. Following Yang & Wang (2012), we classified MATs as either transformation-based tests, regression-based tests, or combination tests.

*Transformation-based tests*

Transformation-based MATs deal with a multivariate problem is by reducing it to a univariate problem, i.e., they create a linear combination of the *m* traits (i.e., the new variate) which can then be used in a univariate analysis. We selected the following transformation-based techniques: sum scores, Principal Component Analysis (PCA), common factor analysis, and the Combined PC test (CPC). Notable, Canonical Correlation Analysis (CCA) is another transformation-based methods. However, assuming an *additive codominant genetic model* in which the GV, coded 0/1/2 for the number of minor alleles, is treated as a continuous predictor (i.e., a "covariate", rather than a "factor"), CCA is identical to MANOVA. In our simulations, we therefore only included MANOVA.

We note that sum score analysis, PCA and factor analysis yield a new variate, which represents all *m* traits in the analysis (see Box 1); this new variate is subsequently used as a dependent variable in a standard univariate regression model in which the GV features as the predictor of interest (in addition to other covariates, e.g., sex, age, covariate correcting for stratification).

*Regression-based tests*

Regression-based MATs centre on the association between the GV and all *m* traits simultaneously. We selected the following regression-based tests: MANOVA, Generalized Estimating Equations (GEE) and MultiPhen. We included two versions of MANOVA: standard MANOVA, which is an *m*-df test (i.e., the individual associations between the GV and the *m* traits are all estimated freely), and a 1-df version of MANOVA in which all *m* regression coefficients of the GV to the *m* traits are constrained to be identical. For GEE, we included 4 versions. First, the background covariance matrix $\mathbf{\Sigma_E}$ (which is modeled as $\mathbf{\Delta_E P_E \Delta_E}$, where $\mathbf{P_E}$ is the residual correlation matrix between the *m* traits conditional on all predictors in the model, and $\mathbf{\Delta_E}$ is a diagonal matrix with the *m* diagonal elements, representing the residual standard deviations of the *m* traits) was either specified as *unstructured* (i.e., all ($m*m-m$)/2 off-diagonal elements in $\mathbf{P_E}$ are estimated freely) or *exchangeable* (i.e., all off-diagonal elements of $\mathbf{P_E}$ are constrained to be identical, i.e., conditional on the *k* predictors, the correlations between the *m* variables are assumed identical, i.e., only 1 parameter is estimated to represent all ($m*m-m$)/2 residual correlations). Second, the regression weights relating the GV to the *m* traits were either all estimated freely (*m*-df test) or constrained to be identical (1-df test).

We did not include Linear Mixed Models (LMM), which are a random effects extension of the basic fixed effects multivariate regression model that MANOVA and GEE are based on, in our main discussion because in the current context, LMM can be written as a GEE exchangeable model without sandwich correction. Because of this similarity, and because sandwich has been shown to be essential when dealing with clustered data (ref), and because LMM often failed to converge in our simulations (especially with *m* > 4), we excluded LMM from our main discussion. Type I and power results are, however, available in the Supplemental Tables.

*Combination tests*

We selected the following combination tests: Nyholt-Šidák and Bonferroni corrected p-values (min-$P_{NS}$, min-$P_{Bonf}$), the Simes test, its adjusted version TATES (Trait-based Association Test that uses Extended Simes), two version of JAMP (Joint genetic Association of Multivariate Phenotypes: $JAMP_{mult}$ and $JAM_{min}$), the meta-analysis inspired techniques $S_{Hom}$ and $S_{Het}$, and the adjusted Fisher-combination test FC-Pearson.

In summary, we considered three types of MATs: transformation-based MATs, regression-based MATs, and combination tests. As an alternative classification, one could distinguish 1-df tests from *m*-df tests. Specifically, 1-df tests either first reduce all *m* traits to a single new variate (transformation-based tests), or constrain all *m* associations between the GV and the *m*-traits to be equal ($S_{Hom}$, and the 1-df versions of GEE and MANOVA). In these tests, the associations between the GV and the *m* traits is modelled by means of a single parameter. In contrast, in *m*-df tests (standard MANOVA, $S_{Het}$, CPC, and the *m*-df versions of GEE), the associations between the GV and the *m* traits are modelled through *m* parameters (allowing variation in the strength of the individual trait-GV associations). As it turned out, this latter classification is more in accordance with the power results: the power results of *m*-df tests were quite distinct from those displayed by 1-df tests, while m-df tests mutually showed very similar power (see Supplemental Information S5 and Table S10-S12), as did 1-df tests (see Supplemental Information S5 and Table S10-S12).

### S3. Alternative classification

In the main text, we chose to follow the conceptual classification of MATs as proposed by Yang and Wang (2012), and distinguished transformation-based MATs, regression-based MATs, and combination tests. Alternatively, one could classify existing MATs based on the statistical behavior of their test statistics. This classification would largely distinguish *m*-df tests, 1-df tests and “minimum-p” test, as described below. The only MAT that does not fit well into this classification is $S_{Het}$, which is mainly due to the fact that the test statistic of $S_{Het}$ is based only on the subsets of traits for which the Wald statistic exceeds a certain threshold, putting it somewhere intermediate between the *m*-df and 1-df tests.

By examining each test statistic, it is in most cases possible to discern the general mathematical structure of the statistic, which can be used to compare and contrast the expected and observed behavior of each test. In practice, it is often useful to abstract away some of the details, in order to simplify the structure of the test statistic and therefore facilitate the comparison of the statistics. As the aim is merely to gain a sense of the general behavior of the methods, this loss of detail does not unduly affect the conclusions that can be drawn.

In particular, most of the test statistics can be decomposed into a kernel $K$ that reflects the relation between the GV and the traits, an inverse residual (co)variance term, and a scaling constant. Only this kernel $K$ is of primary interest. The scaling constants do not affect the functional form of the test statistic at all, and the residual covariance term is typically a function of the kernel and is therefore not crucial for understanding the structure either.

Often, it is also useful to inspect an approximation of the test statistic instead of the test statistic itself, again for ease of comparison. As the results of the simulations bear out, despite such simplifications the comparison that can thus be made match the observed similarities and differences in statistical behavior quite well.

For analyzing the test statistics with sample size $N$ and number of traits $m$, denote the GV as $X$ (a $N \times 1$ vector) and the traits as $Y$ (a $N \times m$ matrix). Except where otherwise specified, it is assumed that both have been centered to have a mean of 0. Consequently, the vector of covariances between the GV and the traits, the key sufficient statistic on which all the test statistics are based, is computed as $S = \frac{Y^T X}{N-1}$. Similarly, the covariance matrix of the traits can be computed as $M_Y = \frac{Y^T Y}{N-1}$

***m-df tests***

For the tests with $m$ degrees of freedom, the (approximate) kernel of the test statistics can be written as a quadratic form, $K = S^T W S$ for some weight matrix $W$.

*Combined Principal Component test (CPC)*

The CPC approach performs a principal components analysis, eigendecomposing the trait covariance matrix $M_Y$ into eigenvector matrix $Q$ and diagonal eigenvalue matrix $\Lambda$: $M_Y = Q\Lambda Q^T$. The traits are then projected onto their standardized principal components $P = YQ\Lambda^{-\frac{1}{2}}$, and the covariances of $P$ and $X$ are computed as $\frac{P^T X}{N-1}$. $\chi^2$ test statistics for these covariances are then computed, summing them to obtain the final test statistic. These $\chi^2$ statistics are highly correlated with the squared covariances however, and as such the kernel of the test statistic is approximately proportional to $(P^T X)^T P^T X = X^T P P^T X = X^T Y Q \Lambda^{-1} Q^T Y^T X = X^T Y M_Y^{-1} Y^T X \propto S^T M_Y^{-1} S$. The weight matrix $W$ for the approximate kernel of the CPC is therefore the inverse covariance matrix of the traits; effectively this compensates for the overlap in the marginal associations contained in the $Y^T X$ vector, with the resulting kernel reflecting the explained variance by $X$ jointly in $Y$.

*MANOVA and Canonical Correlation Analysis*

As noted in Box 1, Canonical Correlation Analysis CCA will under an additive allele model behave the same as MANOVA, and the two are therefore treated as one here. For MANOVA, the test statistic is based on the matrix of model (or between-groups) sums of squares and crossproducts $B$ and the residual (or within-groups) sums of squares and crossproducts $R$, which by definition is equal to $Y^T Y - B$. Different MANOVA test statistics have been developed over the years but they are all based on the eigenvalues of the matrix $BR^{-1}$. The most straightforward is the Lawley-Hotelling trace defined as $tr(BR^{-1})$, which is equal to the sum of the eigenvalues.

To obtain an approximation of this, first we can note that the MANOVA model is equivalent to a multivariate regression model with a predictor matrix $D$ containing three dummy variables encoding the three possible genotypes, such that $Y_i \sim \mathrm{MVN}(\mu^T D_i, \Sigma)$ for each individual $i$, with $\mu$ a $3 \times m$ matrix of group mean parameters and $\Sigma$ the residual covariance matrix. In terms of explained variance, this parameterization is equivalent to using an intercept and only two dummy variables. This can be approximated by assuming a simpler model postulating a linear relation between the group means, replacing the two dummies with the (uncentered) GV. It can in turn be observed that if both $X$ and $Y$ are centered the intercept will be zero and can be removed without changing the model fit, to obtain the model $Y_i \sim \mathrm{MVN}(X_i \beta, \Sigma)$ with $\beta$ is a $m \times 1$ vector of effect sizes. The estimate of $\beta$ is $\frac{Y^T X}{X^T X}$, and hence the matrix $B$ is proportional to $Y^T X (Y^T X)^T \propto SS^T$, and consequently the (Lawley-Hotelling) test statistic is proportional to $tr(BR^{-1}) = tr(SS^T R^{-1}) = tr(S^T R^{-1} S) = S^T R^{-1} S$. In practice, especially under the null hypothesis, $R$ will be dominated by and roughly proportional to $Y^T Y$, and substituting this for $R$ gives a form for the kernel proportional to $S^T M_Y^{-1} S$. Based on this, it would therefore be expected that

MANOVA and CPC will have almost the same power behavior, provided the Type I error rates are correctly controlled.

*Generalized Estimating Equations (GEE)*

In terms of its model, GEE is similar to MANOVA in that it specifies a model of the form $Y_i \sim \mathrm{MVN}(\mu^T D_i, \mathrm{V})$, with $V$ the residual covariance. Test statistics for GEE are harder to approximate however, since GEE is primarily an iterative estimating procedure. It specifies a working residual covariance matrix for $V$, assuming a particular structure as described in Box 2. It then estimates the parameters in the linear model given $V$, then re-estimates $V$ using the estimated parameters. This continues until the estimates converge. As a result the test statistic does not have a closed mathematical form that can be inspected.

What can be noted, however, is that GEE provides consistent estimates of the parameters in the linear model, even if the residual covariance structure is misspecified. Since the linear model parts in MANOVA and GEE are the same, the implied test statistic for GEE is likely to be roughly proportional to that of MANOVA. If the Type I error rates are properly controlled, power for the two methods can therefore be expected to behave in the same way as well.

*MultiPhen*

For MultiPhen an ordinal regression model is used with $X$ as outcome and $Y$ the predictors. Ordinal regression also does not have a convenient closed form. However, as the genotype has only three levels, the fit of the model will be very similar to that of a linear regression model, especially at larger sample sizes. For a linear regression model of the form $X = Y\beta + \varepsilon$, the parameters are estimated as $\hat{\beta} = (Y^TY)^{-1}Y^TX$ and the sum of model squares has the form $\hat{\beta}^T Y^T Y \hat{\beta} = X^TY(Y^TY)^{-1}Y^TY(Y^TY)^{-1}Y^TX = X^TY(Y^TY)^{-1}Y^TX \propto S^T M_Y^{-1} S$. It thus arrives at the same approximate form as CPC and MANOVA, suggesting that these will all behave in much the same way.

*$JAMP_{\mathrm{multi}}$ and FC-Pearson*

For $JAMP_{\mathrm{multi}}$ and FC-Pearson, the test statistic is computed as the sum of -log p-values of the associations of each trait individually with the GV. The -log of a p-value for a correlation test is itself strongly correlated to the squared correlation itself, and as such these test statistics will be proportional to the sum of squared correlations between $X$ and $Y$. With $V_Y$ the diagonal matrix of variances of $Y$, the correlations are proportional to $V_Y^{-\frac{1}{2}}S$ and therefore the test statistic is proportional to $S^T V_Y^{-1} S$. The weight matrix $W$ in this case is therefore equal to $V_Y^{-1}$ rather than $M_Y^{-1}$ as in the other tests. Note that if the correlations between the *m* traits are homogenous (like in our 1-factor scenarios 1-5), these two weight matrices will show the same behavior.

In our current simulations, in which the effect size of the GV was defined on the marginal (i.e., on the level of the individual trait, and not on the level of the set of *m* traits), we see that in general, the power of m-df tests, and thus $JAMP_{\mathrm{multi}}$ and FC-Pearson, decrease with increasing correlations between the *m* traits (scenario 1) because the 'signal-to-noise' ratio decreases (see also Supplemental Information S8 below). Specifically, when the *m* traits correlate more, they share more variance, i.e., the variance common to the traits is larger. Given a fixed effect size of the GV on the level of the individual traits, the relative contribution of the GV to that common variance gets smaller when the traits co-vary more strongly. That is, the signal-to-noise ratio (*signal* referring to the part of the common variance that is due to the (shared) effect(s) of the GV under study; *noise* referring to the remaining part of the common variance that is not due to the GV) is generally better when the covariance between the traits conditional on the GV is low. In the context of $JAMP_{\mathrm{multi}}$, the test statistic $G_o$ (i.e., the sum of the –log10 of the *m* univariate p-values, see Box 3) itself is unaffected by the correlations between the *m* traits, but the distribution of the test statistics as obtained through permutation is not. While other $m$-df tests, which model all $m$ traits simultaneously, can profit from the presence of opposite effects or from unaffected traits in the set (scenarios 2-5; see also Supplemental Information S9), JAMPmulti, based on univariate p-value information only, does not and therefore loses power as a function of the trait correlations in all scenarios.

***1-df tests***

For the 1-df tests, the kernel will be of the form $K = (w^T S)^2$ with $w$ a vector of weights, i.e., the square of a weighted sum of the covariances in $S$. Note that this is equivalent to $S^T w w^T S$, and thus the kernel of the 1-df tests is the same as the kernel of an $m$-df test with $W = ww^T$. The crucial difference is that for 1-df tests $W$ is a matrix with rank of 1, whereas for the $m$-df tests it will have a rank of $m$ (unless $M_Y$ is not of full rank, though in this case it would also not be invertible).

*Sum score, Principcal Component Analysis (PCA, and factor analysis (FA)*

For the sum score, PCA and FA approaches, this form arises from the fact that each of them constructs a new univariate trait, i.e., new variate, $T = Yw$ as a weighted sum of the individual traits, and then test the covariance of this trait with $X$. This covariance is computed as $\frac{T^T X}{X^T X}$, which is proportional to $T^T X = w^T Y^T X \propto w^T S$, the square of which forms the kernel for a $\chi^2$-test or F-test. For the sum score approach $w = \vec{1}$, whereas for PCA and FA $w$ is equal to the eigenvector for the first PC or the factor loadings for the first factor, respectively.

The sum score approach thus gives each trait equal and positive weights in the test, whereas PCA and FA favor traits that load more strongly on the first PC or factor. Notably, in doing so they will also account for the sign of correlations between traits, making it less likely that associations of negatively correlated traits will cancel out against each other in the test statistic.

If the traits are standardized prior to analysis, for each trait $j$ the weight would be adjusted by dividing it by $SD(Y_j)$ (this is equivalent to replacing the covariances in $S$ with corresponding correlations between $X$ and $Y$); for PCA and FA, this is in addition to any changes that result from the analysis of a correlation matrix yielding a different set of eigenvectors and factor loadings.

*1-df MANOVA*

For the 1-df version of MANOVA, the group means are constrained to be the same across the phenotypes. Starting from the simplified model $Y_i \sim \mathrm{MVN}(X_i\beta, \Sigma)$ obtained above, this changes this model to $Y_i \sim \mathrm{MVN}(X_i \vec{1}\beta^*, \Sigma)$, where $\beta^*$ is a scalar reflecting the effect of $X$ on each $Y$. It follows that the mean of the traits $\frac{\vec{1}^T Y_i}{m} \sim \mathrm{MVN}(X_i\beta^*, \sigma^2)$, where $\sigma^2 = \vec{1}^T \Sigma \vec{1}$ is a new residual variance parameter that is the sum of all elements of $\Sigma$. Without loss of information on $\beta^*$, we can therefore substitute this simple regression model for the constrained 1-df MANOVA model. This model is also equivalent to the sum score model, since the scaling by $\frac{1}{m}$ is constant and therefore has no impact on the behaviour of the test statistic.

*1-df Generalized Estimation Equations*

The same principle holds for the 1-df version of the GEE model, reducing it to a univariate regression as well. Since the residual covariance matrix in doing so is compressed to a scalar the specified structure of the residual covariance becomes virtually irrelevant, with the model essentially a simple regression with robust standard error estimation. Since the robust standard errors do not affect the kernel of the test statistic however, the behavior for this model will be the same as the 1-df MANOVA and the sum score analysis.

*$S_{Hom}$*

For the $S_{Hom}$ test, the Wald test statistics of the associations between the GV and each of the traits are used, which are very strongly correlated with the corresponding correlations and thus proportional to $V_Y^{-\frac{1}{2}}S$. These are multiplied by the inverse of trait correlation matrix $R$ and then summed, to obtain $\vec{1}^T R^{-1} V_Y^{-\frac{1}{2}} S = \vec{1}^T V_Y^{\frac{1}{2}} M_Y^{-1} S$. This is then squared in the test statistic. Consequently, this results in $w = M_Y^{-1} V^{\frac{1}{2}} \vec{1}$ as the weight vector. Akin to the *m*-df tests with $W = M_Y^{-1}$, this has the effect of decorrelating the test statistics, assigning less weight to traits that are more strongly correlated with other traits to compensate for the partial duplication of association signals that would propagate across those correlations.

***Minimum p-value tests***

As described in the main text, Table 1 of the main text, and Box 3, combination tests that evaluate the hypothesis that at least 1 of the *m* traits is associated to the GV all constitute some sort of correction for multiple testing, and therefore form a clear third category. In effect, they all have the same test statistic: the minimum p-value across the traits. Differences in the performance of the different methods in this category should therefore entirely be due to differences in ability to control the Type I error rate correctly, with no differences in power beyond this.

### S4. Simulations

*General*

All simulations involved a sample of 2000 unrelated individuals, and one biallelic SNP, which we denoted as the disease susceptibility locus (GV), regardless of whether it was associated with the phenotype(s). The minor allele frequency of this GV was set to .2. We simulated data given *m* standard normally distributed traits (N~(0,1)) that together formed a set of uniformly correlated traits (i.e., a phenotypic 1-factor model) or a set of clustered traits (i.e., a phenotypic 2-factor model). The number of traits *m* was set to 4, 8, or 16. In determining the power, the number of simulations was set to Nsim=1000 per scenario, and we counted the number of times that $p < .05$. To establish the Type I error rates, Nsim was set to 1.000.000, and we counted the number of times that $p < .05$, .01, and .001.

*Phenotypic model*

Conditional on the GV, phenotypic data were *m*-variate standard normally distributed, with the variance-covariance structure corresponding either to a 1-factor model (i.e., uniformly correlated traits), or a 2-factor model (i.e., clusters of traits). In the two factor model, the factors were correlated and each of the *m* phenotypes loaded on only 1 factor (i.e., simple structure, see Supplemental Figure S1 for an illustration with 4 traits). The *m*-variate data were simulated according to the model

$$\mathbf{\Sigma}=\mathbf{\Lambda}*\mathbf{\Psi}*\mathbf{\Lambda}^t+\mathbf{\Theta}. \quad \text{(eq 1)}$$

Here, **Σ** is the *m*x*m* variance-covariance matrix of the *m* traits. In case of a 1-factor model, **Λ** is a *m*x1 matrix of factor loadings with all loadings values set to .32, .55, .77, or .92, which implies correlations between all *m* phenotypes of ~.1, ~.3, ~.6, or ~.85, respectively (i.e., uniformly correlated traits). The 1x1 matrix **Ψ** contains the factor variance, which equals 1 (standard scaling in the common factor model).

In case of a 2-factor model (which we used to model clusters of traits), **Λ** is a *m*x2 matrix, with the first *m*/2 traits loading on the first factor only, and the second *m*/2 traits loading on the second factor (all other loadings fixed to zero). The non-zero factor loadings equaled .55, or .84, so that traits loading on the same factor correlated either ~.30 or ~.70. The matrix **Ψ** is the 2x2 factorial correlation matrix with the variances of the factors set to 1 (standard scaling) and the correlation between factors set to -.9, -.5, -.15, 0, .15, .5, or .9. This implies that correlations between the traits loading on different factors varied from -.27 (.55*-.9*.55) to .27 (.55*.9*.55) for traits correlating .30 within factors, and from -.63 (.84*-.9*.84) to .63 (.84*.9*.84) for traits correlating .70 within factors.

In both 1-factor and 2-factor models, **Θ** is a *m*x*m* diagonal matrix of residual variances. The values of these residual variances were chosen so that the total variance conditional on the GV equaled 1 (e.g., if the factor loading of a variable was set to .32, implying that .32*1*.32=.10 of the trait's variance was explained by the factor, then the residual variance was set to 1-.32*1*.32=.90).

*Genetic data*

A single diallelic GV with minor allele frequency (MAF) of .2 was simulated. The GV affected one, half (*m*/2), or all (*m*) traits directly. The GV explained .1, .2, or .5% of the variance in and individual traits, i.e., on the level of the individual traits. Note that due to the correlations between the *m* traits in a set, the variance explained on the level of the set (i.e., the set of *m* traits) varies across scenarios and within scenarios as a function of the correlations between the *m* traits (see also below). In case of clustered traits (i.e., two-factor model), where half of the observed variables were affected, all these affected traits loaded on the same factor (Supplemental Figure S1).

When the GV was associated to multiple traits (i.e., half or all), the GV either affected all associated traits in the same direction (i.e., all positive effects), or conveyed positive effects to half of the affected traits, and similarly large yet negative effects to the other half (i.e., opposite effects scenario). In scenarios with opposite effects, negative GV-effects were modelled on the "even" traits and positive GV-effects on the "uneven" traits, such that in the 2-factor scenarios, traits loading on the same factor could be positively as well as negatively affected.

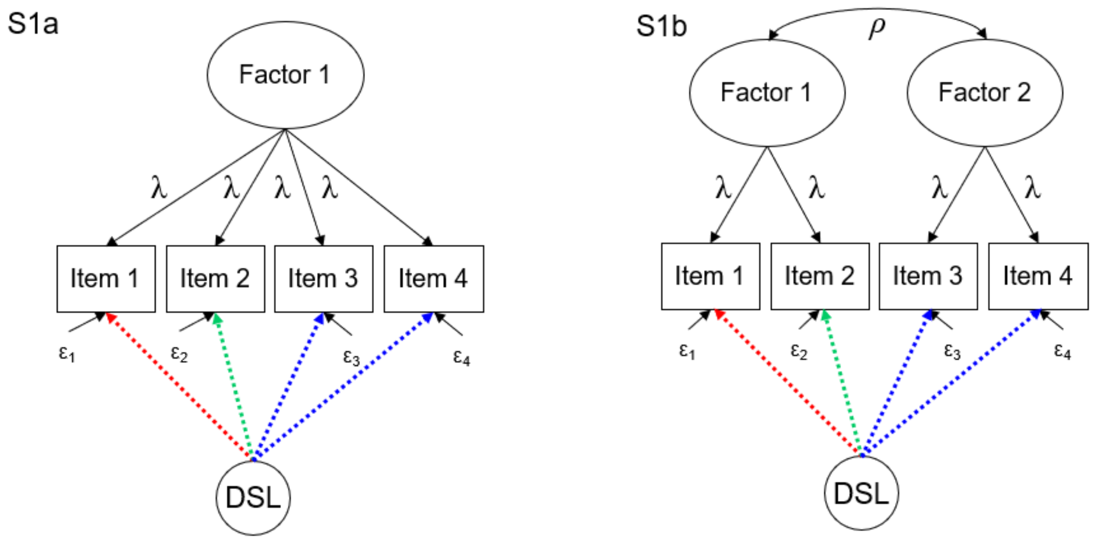

**Figure S1: Schematic representations of the simulated models, illustrated for *m*=4.**
**S1a**: 1-factor model. The factor loadings λ were equal for all traits and were set to .32, .55, .77, or .92, corresponding to correlations between all *m* traits of ~.1, ~.3, ~.6, or ~.85, respectively.
**S1b**: 2-factor model with simple structure, i.e., each trait only loaded on one factor. The factor loadings λ were equal for all traits and were set to either .55 or .84, corresponding to within-factor trait correlations of ~.30 and ~.70, respectively. Factorial correlations *ρ* were set to -.9, -.5, -.15, 0, .15, .5, or .9, so that correlation between traits of different factors ranged from -.27 to .27 for traits correlating .30 within factors, and from -.63 to .63 for traits correlating .70 within factors.

The DSL affected either only one (green), half (blue) or all of the *m* traits. If the DSL affected half of the traits, then all affected traits belonged to the same factor (S1b, blue). If the DSL conveyed opposite effects, then even traits were affected positively (green), and uneven traits negatively (red).

*Simulation settings*

The settings for the 20 Type I error simulation scenarios (Tables S1) and the settings of the 15 power simulation scenarios (Tables S3) are summarized below. Table S2 and S4 summarize 20 Type I and the 15 main power scenarios that we discuss in the main text.

**Table S1 – Overview Type I simulation settings**

| | | | | |
|---|---|---|---|---|
| | ***1-factor*** | | | |
| **# variables** | 4 | 16 | | |
| **Phenotypic correlations** | .1 | .85 | | |
| | ***2-factor*** | | | |
| **# variables** | 4 | | 16 | |
| **Phenotypic correlations within** | .3 | .7 | | |
| **Phenotypic correlations between** | -.9 | -.15 | .15 | .9 |

Note. All Type I simulations were run 1 million times with sample size of N=2000.

**Table S2 – Overview 20 Type I simulation settings**

***Uniform (1-factor)***

| Nvar | Cor | Class |
|---|---|---|
| 4 | 0.1 | 4L |
| 16 | 0.1 | 16L |
| 4 | 0.85 | 4H |
| 16 | 0.85 | 16H |

***Clustered (2-factor)***

| Nvar | Cor within | Cor between | Class |
|---|---|---|---|
| 4 | 0.3 | -0.9 | 4L |
| 16 | 0.3 | -0.9 | 16L |
| 4 | 0.3 | -0.15 | 4L |
| 16 | 0.3 | -0.15 | 16L |
| 4 | 0.3 | 0.15 | 4L |
| 16 | 0.3 | 0.15 | 16L |
| 4 | 0.3 | 0.9 | 4L |
| 16 | 0.3 | 0.9 | 16L |
| 4 | 0.7 | -0.9 | 4H |
| 16 | 0.7 | -0.9 | 16H |
| 4 | 0.7 | -0.15 | 4H |
| 16 | 0.7 | -0.15 | 16H |
| 4 | 0.7 | 0.15 | 4H |
| 16 | 0.7 | 0.15 | 16H |
| 4 | 0.7 | 0.9 | 4H |
| 16 | 0.7 | 0.9 | 16H |

Note. Uniform: *Cor* denotes the correlation between the variables. Clustered: *Cor within* denotes the correlations between variables from the same cluster. *Cor between* denotes the correlation between the 2 clusters. Multiplying *Cor within* by *Cor between* gives the correlation between variables from different clusters.
We used *Class* to create Figure 1 from the main text, in which the Type I error rates were plotted separately for Nvar=4 and Nvar=16, combining scenarios with generally low (L) or generally high (H) trait correlations.

**Table S3 - Overview Power simulation settings**

| | | | | | | | |
|---|---|---|---|---|---|---|---|
| | ***1-factor*** | | | | | | |
| **# variables** | 4 | 8 | 16 | | | | |
| **Phenotypic correlations** | .1 | .3 | .6 | .85 | | | |
| **Affected** | All | Half | 1 | | | | |
| **Effect size** | .01 | .02 | .05 | | | | |
| **Opposite** | True | False | | | | | |
| | ***2-factor*** | | | | | | |
| **# variables** | 4 | 8 | 16 | | | | |
| **Phenotypic correlations within** | .3 | .7 | | | | | |
| **Phenotypic correlations between** | -.9 | -.5 | -.15 | 0 | .15 | .5 | .9 |
| **Affected** | All | Half | 1 | | | | |
| **Effect size** | .01 | .02 | .05 | | | | |
| **Opposite** | True | False | | | | | |

Note. All power simulations were run 1000 times with sample size of N=2000. The effect size is expressed as percentage of variance explained in each affected standardized trait.

**Table S4 – Overview 15 main power simulation scenarios**

| **Scenario** | ***Structure data*** | ***Location GV effect*** | ***Opposite effects*** | ***Corr_within*** |
|---|---|---|---|---|
| **1** | Uniform | All | F | NA |
| **2** | Uniform | All | T | NA |
| **3** | Uniform | Half | F | NA |
| **4** | Uniform | Half | T | NA |
| **5** | Uniform | 1 | NA | NA |
| **6** | Clustered | All | F | .3 |
| **7** | Clustered | All | F | .7 |
| **8** | Clustered | All | T | .3 |
| **9** | Clustered | All | T | .7 |
| **10** | Clustered | Half | F | .3 |
| **11** | Clustered | Half | F | .7 |
| **12** | Clustered | Half | T | .3 |
| **13** | Clustered | Half | T | .7 |
| **14** | Clustered | 1 | NA | NA |
| **15** | Clustered | 1 | NA | NA |

Note. *Structure data*: Uniform refer to data with a phenotypic 1-factor structure and uniformly correlated traits. Clustered refers to a phenotypic 2-factor structure with traits that correlate either .3 or .7 within clusters, while correlation between clusters vary. *Location GV effect*: refers to whether the GV affects 1, half or all the simulated traits. *Opposite effects*: refers to whether the GV affects some traits positively and some negatively (opposite is True: T) or whether the GV-effect has the same sign for all affected traits (Opposite is FALSE: F). *Corr_Within*: describes the values of the correlations between traits belonging to the same cluster (i.e., loading on the same factor). *NA* refer to "Not Applicable".

Important to note with respect to the effect size of the simulated GV:
We defined the effect size on the level of *the affected trait* given the chosen allele frequencies (MAF=.2 in all simulations) and the variance conditional on the GV (which was 1, assuming standard normally distributed traits conditional on the GV). That is, the effect size was defined on the level of the individual variable (i.e., "in the univariate case"), i.e., not accounting for indirect GV-effects (i.e., GV-effects that are picked up by unaffected traits because of their correlation to affected traits (i.e., spillover)). As a result, the power between MATs can be compared within scenarios (where the power of individual MATs fluctuate as a function of the correlations between the traits), but ***not*** across scenarios (because it is, for instance, possible that the total variance explained by the GV in the full set of *m* traits is lower in a scenario where all traits are affected compared to a scenario in which only half of the traits are affected but the correlations between the *m* traits are generally higher, i.e., more spill-over).

*Software*

All simulation were run in R (R Core Team, 2018). For those MATs that were run using dedicated R packages or functions, an overview is given in Table S5. The R-code for the MATs that were fitted in a customary fashion, can be retrieved from the shared simulations scripts (https://ctg.cncr.nl/software/).

**Table S5 – overview of dedicated R-packages used to run MATs**

| ***MAT*** | ***R-packages / R-function*** |
|---|---|
| Standard MANOVA (m df) | manova |
| Customized MANOVA (1 df) | openmx |
| GEE exchangeable m-df | gee |
| GEE exchangeable 1-df | gee |
| GEE unstructured m-df | gee |
| GEE unstructured 1-df | gee |
| MultiPhen | MultiPhen |
| PCA | princomp |
| CPC | princomp + customized code |
| Factor-scores | factanal |
| LMM | lm |

For a subset of MATs for which we believe additional information is required, we provide details on simulation settings and methods below.

*Factor analysis*: We ran a single common factor model in R using the function “factanal”, and calculated factor scores according to the regression method (Lawley and Maxwell, 1971), as implemented in factanal. The factor scores were used in subsequent regression analyses. We note that in principle, one does not need to fit a single common factor model, but could actually use exploratory factor analysis (EFA) to explore the dimensionality of the data. E.g. if EFA would show that the covariance matrix of the trait data can best be described by 3 phenotypic factors, one could calculate factor scores for each of these 3 factors separately, and use the resulting 3 variables in genetic analyses. This way, the factor scores would do more justice to the dimensionality of the data. Here, however, we focus on factor analysis as a data reduction technique to reduce multivariate to univariate data, under the assumption that the single common factor may be viewed as a substantive variable, i.e., as a common cause of the covariance observed in the *m*-variate data.

*PCA*: PCA was run using the R-function “princomp”. To evaluate the association of the GV to the first PC (PC1), the extracted PC1 was saved and used as dependent variable in univariate regression analyses. Like with common factor analysis, one could also use PCA to explore the dimensionality of the data and select not only the first PC but also subsequent PCs for genetic analysis, e.g., when PC1 does not explain a lot of variance, and/or subsequent PCs explain considerable additional amounts of variance, which will be the case when the data shows clustering (multi-dimensional). In that case, selecting multiple PCs for subsequent genetic analysis would do better justice to the dimensionality of the trait data. Here, however, we focus on PCA as a data reduction technique to reduce multivariate to univariate data, under the assumption that PC1, as it explains the most variance, is of most interest.

*CPC*: For the CPC test, all *m* extracted orthogonal PCs were used as dependent variables in a simple univariate regression. Next, the CPC test statistic was obtained by summing the variance explained by the GV in each of the *m* PCs, multiplying this sum by the sample size (N=2000 in our simulations). This test statistic was subsequently evaluated by reference to a $\chi^2$-distribution with *m* degrees of freedom (Aschard et al., 2014).

*MANOVA*: The standard *m*-df MANOVA was evaluated in R using Pillai-Bartlett's test statistic (but note that with one predictor, i.e., the GV, the p-values of the four MANOVA test statistics (i.e., Hotelling-Lawley's trace, Pillai-Bartlett's trace, Wilks' lambda, Roy's largest root) are all identical.
Note that the GV is coded 0/1/2 for the number of minor alleles (additive model) and treated as a covariate , rather than a "factor". In that case, the results obtained using MANOVA are identical to those obtained using Canonical Correlation Analysis, as is built in in PLINK (Ferreira & Purcell, 2009, see also Van der Sluis et al., 2010; Medland & Neale, 2010; Campbell & Taylor, 1996).
For reasons of comparison, we also specified a 1-df MANOVA in the R package Openmx (Neale et al., 2016) by first constraining all *m* associations between the GV and the *m* individual traits to be the same, and then fixing this one parameter to 0.

*GEE*: Generalized estimating equation (GEE) based regression, as implemented in the R library "gee", was used to conduct 1-df and *m*-df tests. GEE is typically used to conduct regression analyses of clustered data, where the statistical model to account for the clustering is misspecified. For instance, the conditional correlation matrix of the traits (conditional on the GV) may be assumed to be diagonal (assuming the traits to correlate 0, i.e., "independence", conditional on the GV), or "exchangeable" (also known as compound symmetric: conditional on the GV all trait correlation are assumed to be equal) while the correlational structure of the traits conditional on the GV is generally more complex in reality. The standard errors are corrected using the sandwich correction (Dobson & Barnett, 2008).

*MultiPhen*: The MultiPhen model was fit using the homonymous R-package "MultiPhen".

*JAMP*: Both versions of JAMP were run using a dedicated R function (see code online). In all our simulations, the number of permutations *J* was set to 1000 for both $JAMP_{mult}$ and $JAMP_{min}$.

$S_{Hom}$: In a meta-analytic fashion, $S_{Hom}$ (Zhu et al., 2015) uses the Wald test statistics obtained in *m* univariate GWA analysses (and possibly across *k* cohorts) to create a new test statistic that follows a $\chi^2$ distribution with 1 df. $S_{Hom}$ accounts for heterogeneity in sample size and for correlations between the test statistics. As a 1-df test, $S_{Hom}$ constraints all GV effects to be the same, and then tests the hypothesis that this 1 GV-parameter is 0. $S_{Hom}$ is thus most powerful when the *m* GV effects are in reality indeed similar in size and sign across the *m* traits.

$S_{Het}$: $S_{Het}$ is equivalent to $S_{Hom}$ but specifically handles heterogeneity in GV-effects across the *m* traits by calculating the new test statistic only for the subset of traits showing a Wald statistic above a certain threshold. This new test statistic is calculated for a range of thresholds, and the maximally obtained value corresponds to $S_{Het}$. The significance of $S_{Het}$ is evaluated against a gamma distribution which can be obtained in 4 ways:

1) Estimate parameters of the gamma distribution based on the correlation matrix observed in the data, and then test the observed $S_{Het}$ value on the gamma distribution derived from the estimated gamma parameters,
2) Estimate parameters of the gamma distribution based on the theoretical correlation matrix (i.e., as used for simulating the data), and then test the observed $S_{Het}$ value on the gamma distribution derived from the estimated gamma parameters,

3) Simulate an empirical gamma distribution based on the correlation matrix observed in the data, and then test the observed $S_{Het}$ on this simulated empirical gamma distribution,
4) Simulate empirical gamma distribution based on the theoretical correlation matrix (i.e., as used for simulating the data), and the test the observed $S_{Het}$ value on this simulated empirical distribution.

Zhu et al (2015) proposed using method 1. However, in a large simulation study like the current, taking the observed correlation matrix for each simulated data is computationally prohibitively slow. We therefore used the theoretical correlation matrix, i.e., the matrix that was used to simulate the multivariate data as this matrix will closely resemble the observed correlation matrix, due to the large sample size (N=2000). Furthermore we used the simulated empirical gamma distribution (method 4), rather than the estimated gamma parameters (method 2), because preliminary simulations showed that the simulated empirical gamma distribution resembles the $S_{Het}$ distribution more closely than the gamma distribution based on the estimated gamma parameters. We thus continued with method 4), but results for method 2) are also shown in Supplemental Tables xxx.

### S5. Type I error rates

*Confidence intervals*

For each of the 20 Type I error rate scenarios (see Table S1), we ran $N_{sim}$=1,000,000 simulations, allowing us to reliably evaluate Type I error rates at α-levels .05 to .001. We note that the large number of simulations provides high statistical power to detect deviations from the expected Type 1 error rate (α), especially for higher α values. Table S6 summarizes the $CI_{99}$ for all α-levels evaluated in our study and clearly shows that the confidence intervals are extremely narrow for the higher α-levels.

**Table S6** 99% Confidence intervals given Nsim=1,000,000 for various levels of α

| α-level | Lower bound | Upper bound |
|---|---|---|
| **.05** | 0.049439 | 0.050561 |
| **.01** | 0.009744 | 0.010256 |
| **.001** | 0.000919 | 0.001081 |

Note. Note that the standard error (SE) of the ML estimator of the p-value is calculated as $SE = \sqrt{\frac{p*(1-p)}{Nsim}}$, where p denotes the percentage of significant tests observed in the simulations (observed Type I error rate given no effect) given the chosen α, and $N_{sim}$ denotes the total number of simulations. The 99% confidence interval for an unbiased nominal p-value when the GV-effect is actually zero thus corresponds to $CI_{99}$=( p -2.576*SE, p +2.576*SE).

*Results study Type I error rates*

We note that the large number of replications ($N_{sim}$=1,000,000 simulations) provides high statistical power to detect small deviations from the expected Type 1 error rate (α), especially for the larger α values. For instance, with 1 million replications, the 99% confidence interval ($CI_{99}$) for α=.05 is very narrow: .04944-.05056 (see Supplemental Table S6 for the $CI_{99}$ for all α-levels). As a result, merely considering which MATs show Type I errors outside the $CI_{99}$ paints a gloomy picture (Supplemental Figure S2a). Type I error rates of MANOVA, $S_{Hom}$, and all transformation-based (i.e., essentially univariate) MATs are virtually always correct. However, when considered across all 20 scenarios and three levels of α (.05, .01, .001, i.e., 60 scenarios in total), all other MATs showed Type I error rates outside the $CI_{99}$, with overall percentages ranging from 22% (CPC) to 92% (FC Pearson) and 100% ($GEE_{uns\_m}$).

However, many of these deviations outside the $CI_{99}$ were very small (see Supplemental Table X). We therefore also looked beyond the $CI_{99}$, by summing the deviations from the expected α across all scenarios within α-levels, allowing us to determine which factors caused the largest deviations (Supplemental Figure S2b). Overall, the largest sums of deviations are observed for Simes, FC-Pearson, $GEE_{uns\_m}$, and min-$P_{Bonf}$. Interestingly, combination tests show mainly deviations from the expected when the *m* traits are highly correlated, while the number of traits *m* mainly drives the deviations in most other method. Taking the direction of the deviations into account (Supplemental Figure S2b), we see that CPC, Simes and min-$P_{Bonf}$ are always conservative, while $S_{Het}$, Tates and min-$P_{NS}$ are conservative when applied to many (highly correlated) traits, and liberal otherwise. All other methods that do show deviations from the expected, always show inflation, with Type I error rates of $GEE_{ex\text{-}m}$, $GEE_{uns\_m}$, and FC-Pearson especially being inflated when *m* is large, irrespective of the correlations between the phenotypes.

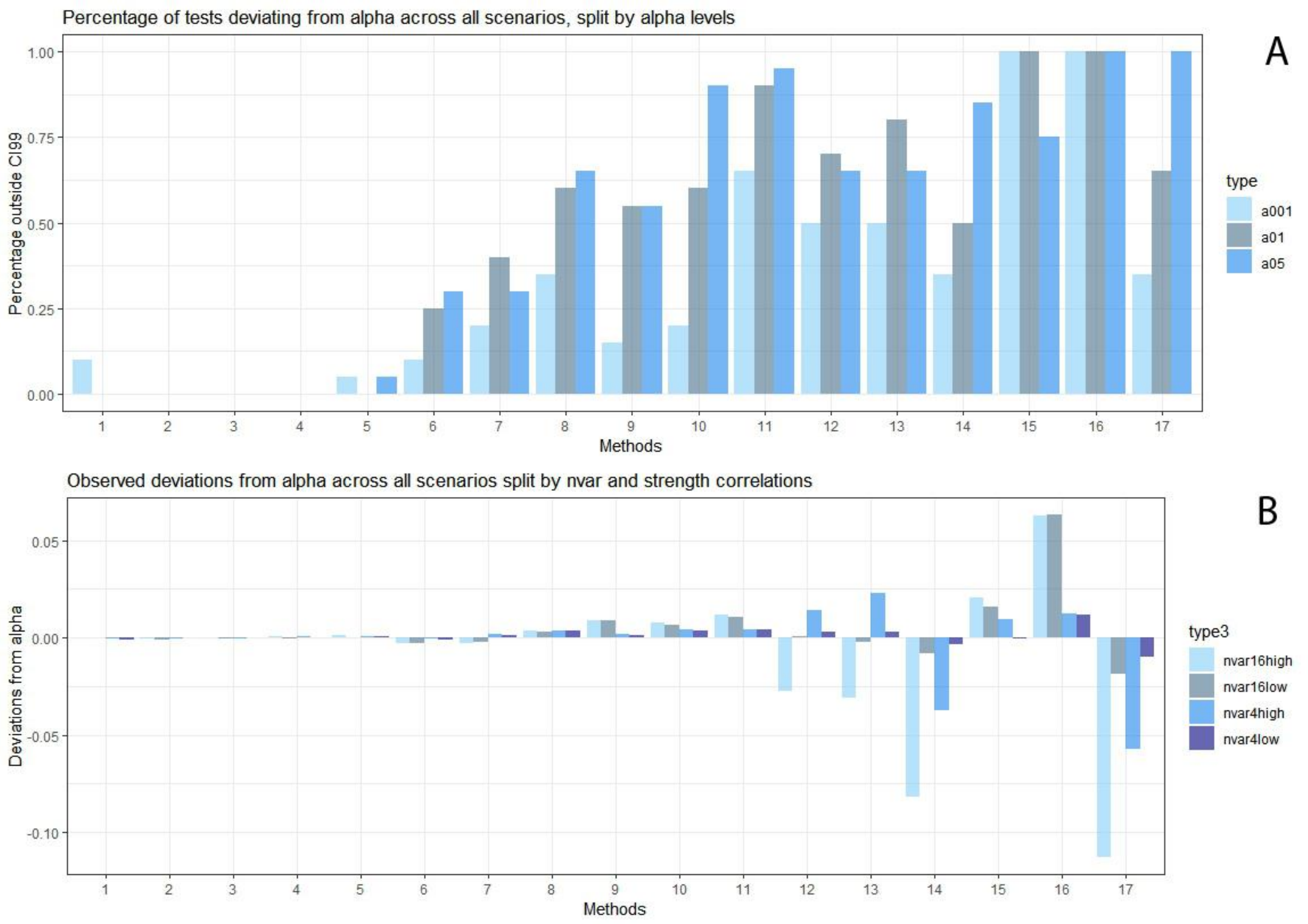

***Supplemental Figure S2: Type I error rates.***
***a****) Across all 20 simulation scenarios, we calculated, separately for 3 levels of alpha (.05, .01, and .001), what percentage of Type I error rates lied outside the 99% confidence interval ($CI_{99}$).*
***b****) Because the $CI_{99}$ is very narrow with Nsim=1,000,000, we also summed, for alpha=.05, the observed deviations from alpha, with larger deviations (in absolute terms) implying larger deviations from the expected value of .05. These deviations can be positive, denoting a liberal test (i.e., too high Type I error rates) or negative, denoting conservative tests (i.e., too low Type I error rates).*
*Methods are numbered: 1=MANOVA, 2=factor score, 3=PCA, 4=sum score, 5=SHom, 6=CPC, 7=SHet, 8=GEEex-1, 9=MultiPhen, 10=MANOVA 1df, 11=GEEun-1, 12=TATES, 13=min-PNS, 14=Simes, 15=FCPearson, 16=GEEun-m, 17=min-PBonf. See Supplemental Tables for Type I error rates given α=.01 and α=.001. Note: the two JAMP-methods were excluded from the Type I error rate study as the correctness of their Type I error rates is guaranteed by their reliance on permutation.*

#### *High Type I error rates explained*

In the main results, the Type I error rates of the following MATs were particularly off: the Type I error rates of the Simes test and the Bonferroni correction (min-$P_{Bonf}$), the Type I error rates of GEE exchangeable and GEE unstructured with *m* df ($GEE_{ex\text{-}m}$ and $GEE_{uns\_m}$), and the FC Pearson test.

*Simes test and the min-$P_{Bonf}$*: Figure 1 in the main text shows that the Simes test and the min-$P_{Bonf}$ are generally too conservative, and specifically conservative when the *m* traits are highly correlated. This is because both methods correct for the *observed number* of traits, which is only correct if the traits are uncorrelated. Because TATES and the Nyholt-Šidák method account (correct) for the correlations among the tests (which are a function of the correlations among the traits), the associated p values fare better in terms of the Type 1 error rate, although here we see an effect of the number of trait, *m*. Specifically, both are slightly liberal when *m* is small, and too conservative when *m* is large.

*FC-Pearson*: The original Fisher Combination test has an increased Type I error rate when the *m* traits are correlated. Given *m* uncorrelated traits, the test statistic is defined as $T = -2\sum_{i=1}^{m} \ln(p_i)$ (Fisher, 1954), which follows a chi-squared distributed with 2*m* dfs. Given *m* correlated traits, it has been shown (Brown and Yang, ref 27/28 in Yang et al, 2016) that, under the null hypothesis of no association between the GV and the *m* traits, *T* follows a scaled chi-squared distribution, or equivalently a specific gamma distribution with shape parameter that can be derived from the mean (μ) and variance ($\sigma^2$) of test statistic T. Yang et al. (2016) established an approximation of μ and σ in case of *m* continuous correlated traits. As our simulation results shown, this approximation is pretty accurate when *m* is small (i.e., *m*=4, as in the original paper by Yang et al, 2016), but when *m* increases to 16, the approximation, and so the Type I error rate, are no longer accurate. The approximation is also less accurate as the correlations among the traits increase (Yang et al., 2016).

*GEE m-df*

Interestingly, while the 1-df GEE variants perform fairly well (quite some p-values outside the $CI_{99}$ range but deviations are minor), the *m*-df variants of GEE (GEE unstructured with *m*-df as shown in the main text, but see paragraph S7 below on that the GEE exchangeable variant with *m*-df yields similar results, as shown in the Supplemental Tables) perform much worse, demonstrating considerable deviations from the expected α, especially when the number of traits *m* is large.
We note that standard GEE software uses sandwich correction of the standard errors of estimated parameters (e.g., of the regression coefficients linking the GV to the *m* traits) to correct for misspecification of $\mathbf{\Sigma}_E$ (Dobson & Barnett, 2008), which almost invariably results from the constraints introduced in $\mathbf{P}_E$ and $\mathbf{\Delta}_E$ (see Box 2). Our results suggest that while the sandwich correction works pretty well when *m* is small, its correction becomes less satisfactory with increasing *m*, i.e., the more parameters are estimated, the more correction is required, and the less accurate the correction is.

#### S6. Power results

In the current study, we conducted power analyses for 18 MATs. However, in the power figures in the main text (Fig X), the power results of various MATs with similar (although not identical, see Supplemental Table XXX) power are represented by a single line. We chose to represent the results of multiple MATs by a single line to increase the clarity of the figures and to avoid clutter. The grouping of the MATs was established through eyeballing of the power results.

Specifically, we first plotted the results for all MATs individually. Next, the results of MATs whose power curves were very similar (i.e., close together and same pattern in all scenarios) were combined and represented by a single line in Figures X. For the simulation scenarios including uniformly correlated traits (i.e., 1-factor models), this resulted in 6 results-groups, i.e., 3 combined groups represented by the results of MANOVA (representing MANOVA, CPC, and MultiPhen), sum scores (representing sum score analysis, PCA, common factor score analysis, $GEE_{ex\_1}$, $GEE_{uns\_1}$, and 1-df MANOVA), and TATES (representing TATES, Simes, $JAMP_{min}$, and min-$P_{NS}$), and 3 lines representing a single MAT, i.e., $JAMP_{mult}$, $S_{Het}$, and the single affected univariate result as a reference. For the simulation scenarios including clustered traits (i.e., 2-factor models), this resulted in 7 results-groups, since for these scenarios PCA and the common factor scores displayed an distinct power pattern.

Important to note: the power results within the TATES-group (representing TATES, Simes, $JAMP_{min}$, and min-$P_{NS}$) were the most heterogeneous: while the methods showed quite similar power and pattern, the deviations between the methods was bigger than was observed between MATs within other groups. TATES was chosen to represent the group as its power was roughly in the middle of the group, with the lines of Simes, min-$P_{NS}$, and $JAMP_{min}$ intertwining with the TATES line across most scenarios.

### S7. Equivalence of GEE *m*-df tests

In this paragraph, we substantiate why the results of GEE *m*-df tests do *not* depend on the choice of working correlation matrix (i.e., “exchangeable” or “unstructured”), while the results of GEE 1-df tests do depend on the choice of working correlation matrix.

**The GEE linear regression model**

Let N be the number of individuals and *m* the number of traits. Let **y** denote the N**m* vector of phenotypic scores, and let **X** denote the N**m* x T design matrix (a.k.a. incidence matrix), which includes the measured disease susceptibility locus (GV), i.e., the predictor of interest. T is the number of parameters to be estimated (including intercepts). Given *m*=4, the design matrix X, as defined by the data of a single subject *i* with genotype $g_i$ (0, 1, or 2) is specified as ($X_i$)

$$\begin{matrix} 1 & g_i & 0 & 0 & 0 \\ 1 & g_i & 1 & 0 & 0 \\ 1 & g_i & 0 & 1 & 0 \\ 1 & g_i & 0 & 0 & 1 \end{matrix}$$

in the case of a 1-df test, and

$$\begin{matrix} 1 & g_i & 0 & 0 & 0 & 0 & 0 & 0 \\ 1 & g_i & 1 & 0 & 0 & g_i & 0 & 0 \\ 1 & g_i & 0 & 1 & 0 & 0 & g_i & 0 \\ 1 & g_i & 0 & 0 & 1 & 0 & 0 & g_i \end{matrix}$$

in the case of a 4-df test (not the only possible parameterization).

The basic regression model is:

$$\mathbf{y} = \mathbf{X\beta} + \boldsymbol{\varepsilon},$$

with residuals $\boldsymbol{\varepsilon}$ (N**m* vector) and $\boldsymbol{\beta}$ a Tx1 matrix of regression parameters. The residuals $\boldsymbol{\varepsilon} = \mathbf{y}-\mathbf{X\beta}$ have covariance matrix $\mathbf{V}_\varepsilon$, which is a N**m* x N**m* block-diagonal matrix, in which the diagonal blocks equal the *m*x*m* covariance matrix of the residuals, i.e., the variances of, and covariances between, the *m* traits conditional on the GV. As we assume homoscedasticity (i.e., conditional on the GV, the variances of all traits are equal, a default assumption in GEE), we can express $\mathbf{V}_\varepsilon$ as

$$\mathbf{V}_{\boldsymbol{\varepsilon}} = \mathbf{I}_\mathbf{N} \otimes \mathbf{S}_{\boldsymbol{\varepsilon}}$$

where $\mathbf{I_N}$ is the NxN identity matrix and $\mathbf{S}_\varepsilon$ is the *m*x*m* homoskedastic covariance matrix. In GEE, $\mathbf{S}_\varepsilon$ equals $s_\varepsilon^2 \otimes \mathbf{r}_\varepsilon$, where $\mathbf{r}_\varepsilon$ is the *m*x*m* residual working correlation matrix, i.e., the correlations between residuals, i.e., between the traits conditional on the GV. The parameter $s_\varepsilon^2$ is the variance of the residuals, i.e., 1 parameter as the variances are assumed equal over traits (i.e., homoscedastic). In conducting the analysis, there are various options for $\mathbf{r}_\varepsilon$: *independence* ($\mathbf{r}_\varepsilon$ is diagonal, residual correlations between traits equal zero), *exchangeable* (all residual correlations between the *m* traits are equal: 1 parameter), or *unstructured* (all residual correlations between the *m* traits are estimated freely, i.e., (*m**(*m*-1))/2 parameters).

The estimate of **β**, denoted **b**, is obtained as follows:

$$\mathbf{b} = (\mathbf{X}^t\mathbf{V}_e^{-1}\mathbf{X})^{-1}\mathbf{X}^t\mathbf{V}_e^{-1}\mathbf{y}$$

where $\mathbf{V}_e$ is the estimate of the covariance matrix of the residuals **e**, which equal

$$\mathbf{e} = \mathbf{y} - \mathbf{Xb}\ .$$

In an iterative procedure, the values of **e** are used to update $\mathbf{V}_e$ (specifically the elements in $\mathbf{r}_e$) which is again used to estimate **b**, etc. By the standard theory (Dobson & Barnett, 2008), we have

$$\mathbf{b} = \left(\mathbf{X}^t\mathbf{V}_e^{-1}\mathbf{X}\right)^{-1}\mathbf{X}^t\mathbf{V}_e^{-1}(\mathbf{X\beta} + \mathbf{e}) = \boldsymbol{\beta} + \left(\mathbf{X}^t\mathbf{V}_e^{-1}\mathbf{X}\right)^{-1}\mathbf{X}^t\mathbf{V}_e^{-1}\mathbf{e}$$

and $\mathbf{V}_b$, the covariance matrix of the estimates in **b**, given the null-hypothesis **β**=0, equals:

$$\mathbf{V}_{b-\text{robust}} = \mathbf{E}[(\mathbf{b} - \boldsymbol{\beta})(\mathbf{b} - \boldsymbol{\beta})^t] = (\mathbf{X}^t\mathbf{V}_e^{-1}\mathbf{X})^{-1}\mathbf{X}^t\mathbf{V}_e^{-1}\mathbf{E}[\mathbf{ee}^t]\mathbf{V}_e^{-1}\mathbf{X}(\mathbf{X}^t\mathbf{V}_e^{-1}\mathbf{X})^{-1}\ .$$

This is the *robust* covariance matrix of the estimates in **b**. The robust standard errors of the estimates in b equal the square root of the diagonal elements. $\mathbf{V}_{b\text{-robust}}$ is robust in the sense that it does not require $E[\mathbf{ee}^t]=\mathbf{V}_e$ , i.e., the working correlation matrix $\mathbf{r}_e$, on which $\mathbf{V}_e$ is based, may be misspecified. Consequently, the covariance matrix of the residuals $\mathbf{V}_e$ does not equal the expected covariances between the residuals $E[\mathbf{ee}^t]$. Conversely, a correctly specified $V_e$, i.e., $E[\mathbf{ee}^t]=\mathbf{V}_e$, implies

$$\mathbf{V}_b = (\mathbf{X}^t\mathbf{V}_e^{-1}\mathbf{X})^{-1}\mathbf{X}^t\mathbf{V}_e^{-1}\mathbf{V}_e\mathbf{V}_e^{-1}\mathbf{X}(\mathbf{X}^t\mathbf{V}_e^{-1}\mathbf{X})^{-1} = (\mathbf{X}^t\mathbf{V}_e^{-1}\mathbf{X})^{-1},$$

the standard expression for the covariance matrix $\mathbf{V}_b$.

As mentioned, GEE regression analysis requires a choice of working correlation matrix $\mathbf{r}_\varepsilon$. Suppose we compare *exchangeable* and *unstructured*. Let the associated matrices $\mathbf{V}_e$ be $\mathbf{V}_{e1}$ and $\mathbf{V}_{e2}$, and let the associated matrix $\mathbf{V}_b$ be $\mathbf{V}_{b1}$ and $\mathbf{V}_{b2}$, such that the estimates of the matrices of regression parameters $\mathbf{b}_1$ and $\mathbf{b}_2$ equal

$$\mathbf{b_1} = (\mathbf{X}^t\mathbf{V}_{e1}^{-1}\mathbf{X})^{-1}\mathbf{X}^t\mathbf{V}_{e1}^{-1}\mathbf{y}$$

and

$$\mathbf{b_2} = (\mathbf{X}^t\mathbf{V}_{e2}^{-1}\mathbf{X})^{-1}\mathbf{X}^t\mathbf{V}_{e2}^{-1}\mathbf{y}.$$

If $\mathbf{b_1}$ equals $\mathbf{b_2}$, the robust standard errors as estimated in $\mathbf{V}_b$ will be equal. This is because then

$$(\mathbf{X}^t\mathbf{V}_{\mathbf{e1}}^{-1}\mathbf{X})^{-1}\mathbf{X}^t\mathbf{V}_{\mathbf{e1}}^{-1} = (\mathbf{X}^t\mathbf{V}_{\mathbf{e2}}^{-1}\mathbf{X})^{-1}\mathbf{X}^t\mathbf{V}_{\mathbf{e2}}^{-1} .$$

Consequently, differences between $\mathbf{V}_{\mathbf{b1\text{-}robust}}$ and $\mathbf{V}_{\mathbf{b2\text{-}robust}}$ arise only if the residuals **e** differ, or, more specifically, if the estimates in $\mathbf{b_1}$ and $\mathbf{b_2}$ differ, i.e., $\mathbf{y}\text{-}\mathbf{Xb_1} \neq \mathbf{y}\text{-}\mathbf{Xb_2}$. Given the *m*-df test, the parameter estimates in $\mathbf{b}_1$ and $\mathbf{b}_2$ are found to be equal, because each trait is regressed on the GV with a unique intercept and slope (in a correctly specified genetic model: model is saturated with respect to the GV). Hence, in this case, we see no difference in standard errors of the parameters in **b** as a function of the chosen working correlation matrix.

### S8. The variance of sum scores

To understand the workings and power of the sum score method, it is useful to consider the variance of the sum of *m* trait scores. Specifically, the variance of the sum of *m* traits is equal to the sum of all the entries of the *m*x*m* variance-covariance matrix between these *m* traits. For two traits $P_1$ and $P_2$

$$\sigma^2_{(P_1+P_2)} = \sigma^2_{P_1} + \sigma^2_{P_2} + 2 * \sigma_{P_1,P_2} , \quad [1]$$

and formulated generally for *m* traits:

$$\sigma^2_{(\sum_{i=1}^{N} P_i)} = \sum_{i=1}^{N} \sum_{j=1}^{N} \sigma_{P_i,P_j} , \quad [2]$$

where $\sigma^2$ and σ denote the variance and covariance, respectively.

Given a set of *m* traits (with fixed variances), the sum of highly positively correlated traits will thus have a larger variance than the sum of less strongly correlated traits because each covariance contributes twice to the variance of the sum. Also, the presence of negatively correlated traits decreases the variance of the sum.

In the context of genome-wide association studies using sum scores, the question is how much the GV of interest contributes to the total variance of the sum of *m* traits. This depends not only on the size of the effects of the GV on the traits, but also on a) how many of the *m* summed traits are affected by the GV, and b) how strongly the *m* traits correlate conditional on the GV.

Summarizing, the variance of the sum, which summarizes the communality between the traits, increases with the correlation among the *m* traits. This implies that given a fixed effect size of the GV on the affected traits in the set of *m* traits, the relative contribution of the GV to that common variance gets smaller when the *m* traits co-vary more strongly. This means that the *signal-to-noise ratio* (*signal* referring to the part of the variance of the sum that is due to the (shared) effect(s) of the GV under study; *noise* referring to the remaining part of the variance of the sum that is not due to the GV) is generally better when the covariance between the traits conditional on the GV is low.
Let us consider some specific scenarios.

*Local GV – affecting only 1 of the m traits*: If a GV affects only 1 of the *m* traits, then the GV only contributes to the variance of the sum through 1 entry of the *m*x*m* variance-covariance matrix, namely through the variance of that 1 affected trait. It is clear that the signal of that GV will contribute little to the variance of the sum, especially if *m* is large and the covariances between the *m* traits are large. That is, the larger the number of traits *m* (of which *m*-1 traits are unaffected!), the larger the *m*x*m* variance-covariance matrix, and thus the more the variance of the sum consists of "noise" (i.e., variance not due to the GV of interest), i.e., the lower the signal-to-noise ratio. So when the GV affects only 1 trait, almost all variances and all the covariances that are summed to make up the variance of the sum contain only noise: the larger the covariances, the more noise is added.

*Local GV –affecting 2 of the m traits*: A GV that affects 2 of the *m* summed traits contributes not only to the variances of both traits but that GV also causes, or contributes to, the covariance between this pair of traits. This GV thus contributes to the variance of the sum of *m* traits in 4 ways: through the variance of trait 1, through the variance of trait 2, and twice through the covariance between the two traits. Like with GVs affecting only 1 of the *m* traits, the signal-to-noise ratio for this GV will be better if a) *m* is small (i.e., the fewer unaffected traits are included in the sum, the better), and b) the covariances between the *m* traits are small.

*Global GV – affecting all m traits*: If a GV affects all *m* traits, then the GV contributes to the sum of the variance of the sum of the *m* traits through every entry of the *m*x*m* variance-covariance matrix. Transformation-based techniques like sum scores therefore can have excellent power to detect global GVs. Given a fixed effect size of the GV, the power of the sum score approach does, however, decrease with increasing covariance between the *m* traits, i.e., if the covariances are large and consist mainly of noise (i.e., covariance not due to the GV under study), then the majority of the variance of the sum is not due to the specific GV, decreasing the signal-to-noise ratio. This is what we see in scenario 1: the power to detect the GV affecting all *m* traits decreases when the correlations between the m traits increase.

*Presence of negatively correlated variables*: Noteworthy, in the calculation of sum scores, the presence of (unaffected) negatively correlated traits can have a beneficial effect on the detection of GV-effects (as we see in scenarios 6-11): the negative covariances between pairs of traits reduce the total variance of the sum, which then improves the signal-to-noise ratio. While the sum score, PCA and factors scores perform very similarly when traits are homogeneously correlated (e.g., scenarios 1-5), their power to detect GV effects is very different when the traits show a clustered nature and the clusters correlate negatively (scenarios 6-11). This is indeed because the power of the sum score actually profits from the presence of negatively correlated traits (variance of the sum decreases, as a result of which the signal-to-noise ratio increases), while the first PC from PCA analysis and the common factor from factor analysis do in this clustered setting not describe the communality of all *m* traits well.

### S9. MANOVA

In this paragraph we explain for two traits (based on Cole et al, 1994), why the power of MANOVA depends not only on the sign and size of the GV effect, but also on the sign of the correlation between the modelled traits.

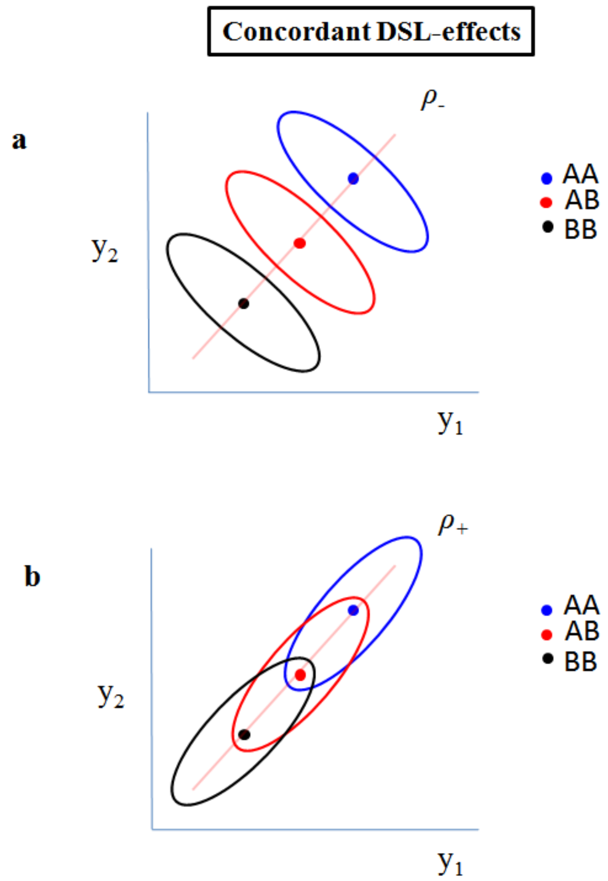

**Concordant GV-effects**

An independent variable, in this case a GV with three levels (i.e., genotypes: AA, AB, BB), affects both variables Y1 and Y2 in the same direction, i.e., the more A-alleles one carries, the higher the scores on both Y1 and Y2. As the GV determines the group membership, the effect of the GV is visible in the means of the three genotype groups (as shown by the position of the centroid).

In the figures, we see three bivariate distributions for genotype groups BB, AB and AA. The group means differ on both variables Y1 and Y2 (i.e., the group-membership variable, which is the GV, has an effect on both variables): group BB scores lowest on both variables (as shown by the position of the centroid), while group AA scores highest on both variables. Ellipses represent the 95% CIs around the centroids.

In the panel **a**, Y1 and Y2 correlate negatively ($\rho$-), and in the panel **b** they correlate positively ($\rho$+). As a result, the bivariate distributions of the genotype group scores either do (panel **b**), or do not (panel **a**), overlap. The extent of overlap between the bivariate distributions determines how well MANOVA will be able to distinguish between the 3 genotype groups using 1 linear combination of Y1 and Y2.

The power to detect a GV that affects both Y1 and Y2 in the same direction, will thus be higher if Y1 and Y2 are negatively correlated.

**Discordant / opposite GV-effects**

A GV affects both variables Y1 and Y2, but in opposite directions (i.e., opposite effects), i.e., the more A-alleles one carries, the higher the scores on both Y2 but the lower the scores on Y1.

In panel **c**, Y1 and Y2 are negatively correlated (*ρ*-): the combination of negative correlations between Y1 and Y2 and opposite effects of the GV causes overlap of the bivariate distributions of the three genotype groups, making it more difficult for MANOVA to distinguish the three groups.

In the panel **d**, the variables Y1 and Y2 are positively correlated (*ρ*+): the combination of positive correlations between Y1 and Y1 and opposite effects of the GV causes minimal overlap between the bivariate distributions of the three genotype groups, i.e., excellent distinction of the three groups.

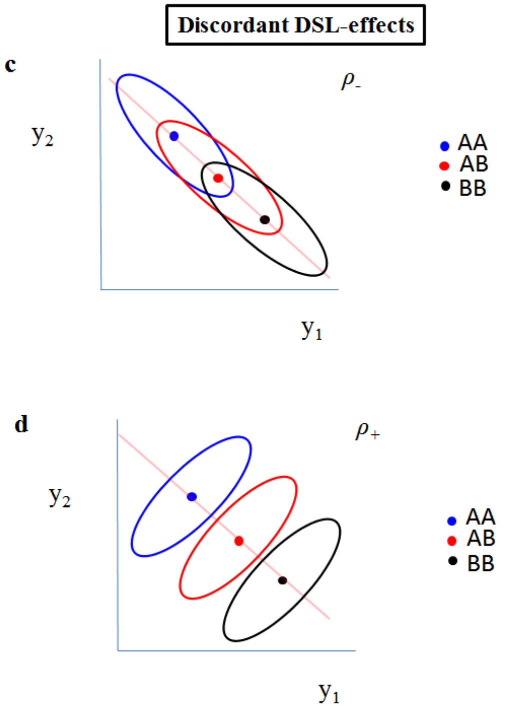

**Selective GV effects**

A GV affecting only one variable, in this case Y2 but not Y1, i.e., the more A-alleles one carries, the higher the scores on Y2, but genotype does not affect the scores on Y1.
In this particular case, the correlation between Y1 and Y2 (negative in panel **e**, positive in panel **f**) does not affect how well the three bivariate distributions of the three genotype groups can be distinguished.

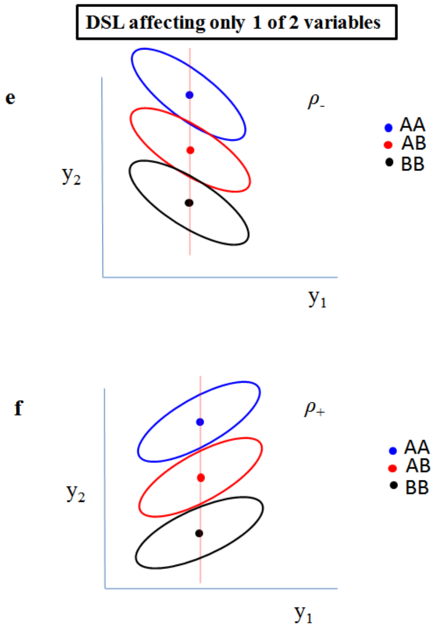

**Summary**

In summary, the critical consideration is not simply the sign (and magnitude) of the correlation coefficient between the traits Y1 and Y2, but the relationship between the sign of the correlation and the pattern of group differences caused by the GV under study (Cole et al. 1994, p 468), i.e., the relation between the sign of the trait correlation and whether/how the GV affects Y1 and Y2.

#### S10. Adding variables: good or bad idea?

Looking at the power results summarized in Figure X of the main text, the power to detect a GV is very often higher when m=16 than when m=4. Based on this, one could wonder whether adding traits to the analysis is advantageous for the power to detect GVs. Here, we want to elaborate on this question from the perspective of MANOVA, because we have seen that MANOVA profits from the presence of traits that are unrelated to the GV, or that are oppositely related to the GV.

First, suppose one disposes of a set of *m* indicators of trait X. One can then add 1 to *m* indicators of X to the multivariate analysis to detect a GV for trait X. If these *m* indicators can indeed all be considered measures of X, then adding them to the multivariate analysis is always wise because:

1) If the newly added indicators are not related to the GV, then the power of MANOVA generally increases because adding unrelated traits increases the power of MANOVA (see also Cole et al., 1994).
2) If the newly added indicators are related to the GV but in an fashion opposite to the relations that the already included indicators have to the GV (i.e., opposite effects), then the power of MANOVA to detect the GV increases (see also Cole et al., 1994)
3) If the newly added indicators are also related to the GV and in the same fashion (i.e., same direction of effect), then the power of MANOVA will decrease, but generally no more than ~10-15% (which is a lot, but the power gain expected in scenarios 1 and 2 described above can be much more extensive).

These three "rules" hold when the *m* indicators can indeed all be considered measures of trait X. However, if the new variables cannot reasonably be considered true indicators (i.e., good measures) of the trait of interest X (e.g., they are phenotypically correlated to variables measuring X, but are not themselves indicators of X, e.g., original set of 5 measures of cognitive ability and adding 1 measure of SES, which is phenotypically correlated to cognitive ability but not actually a measure of cognitive ability), then the problem arises that this new variable can add its own idiosyncratic genetic signal that may or may not be genetic signal for trait X. If this idiosyncratic signal is not shared with X, then this signal can thus be considered false positive signal for the original set of measures, and this false positive signal for X (i.e., genetic signal that is not biologically informative for the original trait X under study).

Overall, results of multivariate analyses will often require follow-up analyses to answer the question which of the *m* traits show association with the GV, or whether the GV effect is shared by all/many of the traits.
